# Too few, too many, or just right? Optimizing sample sizes for population-level inferences in animal tracking projects

**DOI:** 10.1101/2025.07.30.667390

**Authors:** Inês Silva, Christen H. Fleming, Michael J. Noonan, William F. Fagan, Justin M. Calabrese

**Affiliations:** Center for Advanced Systems Understanding (CASUS), Helmholtz-Zentrum Dresden-Rossendorf (HZDR), Görlitz, Germany; Department of Biology, University of Central Florida, Orlando, Florida, USA; Smithsonian National Zoo and Conservation Biology Institute, Virginia, USA; Department of Biology, University of British Columbia Okanagan, Kelowna, Canada; Okanagan Institute for Biodiversity, Resilience, and Ecosystem Services, University of British Columbia Okanagan, Kelowna, Canada; Department of Computer Science, Math, Physics, and Statistics, University of British Columbia Okanagan, Kelowna, Canada; Department of Biology, University of Maryland, Maryland, USA; Department of Ecological Modelling, Helmholtz Centre for Environmental Research — UFZ, Leipzig, Germany

**Keywords:** movement ecology, experimental design, home range, space use, movement behavior

## Abstract

1. Successful animal tracking projects depend on well-informed sampling strategies and robust methods to yield biologically meaningful inferences. Considering financial and logistical constraints, the reliability of research outputs is shaped by key decisions regarding study duration (*how long* should each individual be tracked?), sampling frequency (*how often* should new locations be collected?), and *how many* individuals should be tracked. To maximize their conservation value, studies must consider estimator precision and avoid biased inferences of key parameters related to movement behavior and space use, as this can lead to wasted resources and misguide management actions.
2. To address these challenges, we propose a workflow for determining the optimal sample sizes for population-level home range area and speed estimates, explicitly addressing the trade-offs between sampling duration (*T*), sampling interval (reciprocal of frequency; Δ*t*), and *population* sample size (*m*). While *a priori* study design is considered best practice, this workflow can be applied at multiple stages, including concurrent with data collection, or as a *post hoc* evaluation.
3. By selecting robust methods that are sampling-insensitive, and by quantifying and propagating uncertainty through downstream analyses, we can determine whether our sample sizes (both at the individual- and population-level) are sufficient to yield robust population-level inferences. Furthermore, researchers can integrate additional logistical constraints such as fix success rate, location error, and potential device malfunctions, while also accounting for individual variation. We illustrate potential applications of this workflow through empirically-guided simulations.
4. To facilitate its use and implementation, we incorporated this workflow into the user-friendly ‘movedesign’ R Shiny application. This application enables researchers to easily test different sampling strategies, and as of version 0.3.3, integrates population-level analytical targets. This workflow has the potential to improve the rigor and reliability of animal tracking projects conducted under logistical and financial constraints, and thereby support more effective scientific research, wildlife management, and conservation efforts. It can also support grant or permit submissions by generating evidence-based recommendations for the number of biologgers needed or sampling effort required to achieve research objectives.

## Introduction

Animal movement underlies and links several critical processes, which scale from individuals to populations and from communities to ecosystems (Hooten et al., 2017). Biologging and tracking technologies are vital for gathering such data, and provide insights into a wide range of topics, including space use (Fleming & Calabrese, 2017; Horne et al., 2019), behavioral niche specialization (Hertel et al., 2021), collective behavior and social networks (He et al., 2022), encounter processes (Martinez-Garcia et al., 2020), and migration routes (Kumar et al., 2020). Home range areas are key metrics in conservation planning, informing the delineation or evaluation of protected areas, and the design of management interventions (e.g., Pekarsky et al., 2021; Schofield et al., 2013). Similarly, changes in movement speed and distance travelled can highlight how animals alter their behavior in response to anthropogenic factors (Doherty et al., 2021), providing valuable insights into energy expenditure, foraging behaviors, or the mitigation of human-wildlife conflict. Although both classes of metrics are useful, the inherent tradeoff between sampling duration and sampling interval often prevents their concurrent estimation with the same study design: questions involving home ranges require longer study durations to capture potential changes in space use, whereas questions that focus on movement speed require shorter sampling intervals to facilitate improved estimates of this metric. Population- and species-level parameters are also essential for advancing ecological understanding and conservation practice. Accurately estimating mean home range areas (while incorporating individual variation and uncertainty) provides a fundamental metric of species’ spatial requirements. These metrics underpin numerous studies that examine broad-scale responses to environmental pressures, such as human disturbance, habitat fragmentation, and climate change (e.g., Benitez et al., 2022; Broekman et al., 2024).

Suboptimal study design is a major underlying cause of reduced inferential potential in ecological research (Purgar et al., 2022), emphasizing the need for workflows that support robust study planning. Effectively integrating movement ecology into evidence-based conservation requires both well-designed sampling strategies and robust analytical methods that generate reliable outputs while mitigating common biases (Fleming et al., 2022; Hayward et al., 2015; Silva et al., 2022; B. K. Williams & Brown, 2019). Determining the optimal number of individuals to track is a major challenge during study design, shaped by funding, logistical and ethical constraints that not only impact the number of individuals included in each study, but the duration and frequency of data collection as well. Relying solely on movement parameters estimated from a single individual is often insufficient to capture the complexity of population patterns. Instead, we need robust population-level estimates that reflect variation and behavior at the population scale, which requires an optimal balance between individual-level sampling parameters (duration and interval) and the total number of tracked individuals. Achieving this balance is a multifaceted optimization problem, heavily dependent on which research questions are being addressed. Ideally, researchers would track as many animals as possible, for as long as possible, and as frequently as possible. However, due to logistical and financial constraints, this level of data collection is often unfeasible. Consequently, researchers are forced to make compromises: track fewer individuals, or for shorter durations, or at coarser resolutions. This is especially important when preparing grant proposals or applying for permits, as researchers must objectively justify the number of biologgers they plan to buy or the number of animals to be captured.

The cost of a single GPS unit can reach several thousands of dollars (Lahoz-Monfort & Magrath, 2021; Spiegel et al., 2022), and each additional tracked individual can incur higher logistical overheads, related to personnel time or field visits (Soanes et al., 2013), as well as the transmission of data feeds. Furthermore, animal welfare guidelines emphasize that researchers should, whenever possible, use lighter and less invasive tags or track only as many individuals as needed (Portugal & White, 2018; Williams et al., 2020). Biologger deployments may have deleterious impacts on breeding productivity, offspring quality, energy expenditure, or survival rates (White et al., 2013). Over time, these stressors can influence population dynamics and exacerbate threats. Determining the number of individuals to track is a critical design decision, and we should aim to balance information yield while limiting impacts on individual fitness. More proactive approaches, through data-informed protocols or simulation-based assessments, can ensure we derive biologically meaningful inferences (Guillera-Arroita & Lahoz-Monfort, 2012; Kaur et al., 2024; Silva et al., 2023). However, both limited sample sizes and the intrinsic behavioral variability within populations increase the risk of producing biased or unreliable estimates (Hertel et al., 2021). Key metrics such as home range areas may be significantly underestimated with conventional methods (Noonan, Tucker, et al., 2019), while inter-individual differences may drive variation in space use and exposure to risk (such as when younger individuals exhibit more exploratory behavior; Forrest et al., 2024). Failure to capture this uncertainty or variability could result in wildlife management and conservation actions based on biased or incomplete information.

This manuscript builds on the study design framework introduced in Silva *et al*. (2023), presenting a comprehensive workflow for identifying optimal sample sizes and expanding it to population-level inferences of mean home range area and mean movement speed. Here, we focus on mean estimates within a single sampled population (hereafter referred to as *population* sample size, *m*), though the workflow can be readily adapted for comparisons across groups (*e.g.*, sex, age, habitat). Researchers can apply this workflow to assess a predefined *population* sample size, or the minimum *population* sample size required to achieve a desired level of estimator precision. Throughout, we employ methods that retain statistical precision, account for common sources of bias, and propagate uncertainty through the use of the continuous-time movement modeling framework (Calabrese et al., 2016; Fleming et al., 2022). This workflow can be executed at various stages to: 1) justify resource allocation in funding proposals and permit applications, 2) refine study design using pilot data *before* data collection, 3) assess the reliability of emerging results *during* data collection as an iterative process (while exercising caution, since modifying study designs based on within-study precision may affect statistical inferences; Kairalla et al., 2012), or 4) assess any findings *a posteriori* and inform future research. The latest version (`v0.3.3`) of the `‘movedesign’ R` Shiny application now fully supports the analysis of population sizes in the context of movement study design. We demonstrate the implementation of this workflow in more detail with a case study involving African buffalos (*Syncerus caffer;* Supplementary File S1), and provide a step-by-step guide using the `‘movedesign’` application (Supplementary File S2). Ultimately, we seek to empower researchers, decision makers, and conservation practitioners to enhance the reliability of animal tracking projects, thereby contributing to both scientific rigor and conservation efforts.

## Workflow

### Overview

To determine optimal sample sizes (both at the individual- and at the population-level), we first must highlight several key concepts. First, without quantifying uncertainty, it will be difficult to distinguish true biological patterns from statistical artifacts or biases (Hallman & Robinson, 2020; Noonan, Tucker, et al., 2019; Silva et al., 2022). Conventional home range estimation methods, such as Minimum Convex Polygons (MCPs) and Kernel Density Estimators (KDEs), remain prevalent even though they lack accompanying confidence intervals and implicitly assume successive locations are independent, which can lead to biases when data are autocorrelated, which is increasingly the norm in high-resolution tracking data. As autocorrelation is a fundamental property of modern movement data, the independence assumption is often violated (Noonan et al., 2019). Conversely, with continuous-time movement models, autocorrelation is reconceptualized as a central and informative characteristic of the movement process (Calabrese et al., 2016; Fleming et al., 2017). As animal movement paths represent realizations of these processes, we can characterize movement behavior directly through autocorrelation timescales: the position autocorrelation timescale, *τ*_*p*_, which reflects how long an animal takes to cross its home range (*home range crossing time*), and the velocity autocorrelation timescale, *τ*_*v*_, which reflects how long an animal persists in its motion (*directional persistence*). These parameters, in turn, impose constraints on the sampling design that must be met in order to capture small-scale and large-scale phenomena (Silva et al., 2023). To understand how, we need to consider the *absolute* sample size (*n*, total number of locations recorded) and the *effective* sample size (*N*, the number of independent and identically distributed locations required to produce the same quality estimate; Fleming et al., 2019). The *effective* sample size (*N*) provides a more reliable estimate of information content of a dataset than either absolute sample size (*n*), sampling duration, or interval separately— and ultimately determines both bias and sampling variance of estimators (Noonan et al., 2019; Silva et al., 2022). For home range areas, the effective sample size (*N*_*area*_) is defined with respect to the position autocorrelation timescale parameter (*τ*_*p*_). Increasing the total number of locations without increasing the total sampling duration, *T*, relative to *τ*_*p*_ does not substantially increase the effective sample size for area. In other words, the effective number of independent locations for home range estimation is determined by the duration of tracking relative to *τ*_*p*_, not by the total number of locations recorded. For example, if an animal typically crosses its home range once per day (*τ*_*p*_ = 1 day), a 10-day tracking period provides approximately the same information as 10 independent locations (*N* ≈ 10), regardless of whether locations are recorded every second, minute, or hour. For movement speed, the effective sample size (*N*_*speed*_) is defined with respect to the velocity autocorrelation timescale parameter (*τ*_*v*_).

Using empirically-guided simulations conditioned on fitted movement models from existing movement datasets, researchers can reproduce movement with the same autocorrelation timescale parameters and strategically evaluate which sampling parameters (duration, *T*, and interval, Δ*t*) and *population* sample sizes (*m*) will adequately capture both individual- and population-level estimates of home range area and movement speed. Therefore, to assess the impact of study design through this simulation-based approach, we recommend five key steps: (i) define research targets; (ii) estimate pilot parameters from available data, (iii) simulate synthetic data based on estimated pilot parameters over a range of study designs, (iv) estimate individual- and population-level parameters for each study design, and (v) assess the degree to which each study design supports robust inference of the population parameter(s) of interest.

### Defining research targets

First, any broad research questions (*e.g.*, “I want to understand space-use patterns of my focal species”) must be translated into specific, quantifiable objectives. This task is crucial, as researchers cannot properly tackle an ill-posed problem (Game et al., 2013), as ill-defined questions lead to ill-defined analyses, which in turn produce unreliable inferences (Gould et al., 2025; Knief & Forstmeier, 2025). Clear objectives reduce these risks by explicitly identifying what estimates the sampling strategy must be designed to support and why. If we are interested in estimating large-scale processes, such as long-term space-use requirements (hereafter, home range), sampling duration ought to be maximized in relation to the *home range crossing time* (*T* ≫ *τ*_*p*_) to detect a sufficient number of home range crossing events (Fleming et al., 2015; Silva et al., 2022). For fine-scale processes, we need to assess which sampling interval can effectively capture the fine-scale processes of interest: for example, anthropogenic effects through changes in mean movement speed (Noonan et al., 2019; Thompson et al., 2021). Speed and distance estimates benefit from shorter sampling intervals (Δ*t*) in relation to directional persistence *τ*_*v*_ (Noonan, Fleming, et al., 2019). For Δ*t* < *τ*_*v*_, *N*_*speed*_ generally increases with sampling duration (*T*). When Δ*t* > 3 τ_*v*_, the data no longer retain a statistically detectable signature of the animal’s directional persistence. Once we establish our research targets, the next step is to select the appropriate datasets to extract individual- and population-level parameters.

### Extracting pilot parameters

Evaluating whether research objectives are achievable requires prior knowledge of the characteristic movement scales of the focal species (Gurarie & Ovaskainen, 2011; Silva et al., 2023). These parameters can be extracted from existing movement datasets, preferably from the same species. If data for the focal species are unavailable, data from a closely-related species with comparable movement behaviors can provide preliminary guidance for designing a pilot study —for example, leopard (*Panthera pardus*) and jaguar (*Panthera onca*), which have similar home range crossing times and behaviors. Movebank currently hosts data for 1,400 species across nearly 8,500 studies, totaling over 6.9 billion locations (www.movebank.org). We encourage researchers to explore this resource, or similar data-sharing platforms, to inform their animal tracking projects. Reaching out to researchers who have previously studied your focal species can provide access to data that may not be publicly available.

Parameter extraction is a crucial step in our workflow, shaping all subsequent simulations (Fig. 1). Therefore, researchers must preprocess the data by removing biologically implausible outliers (Gupte et al., 2022; Noonan, Fleming, et al., 2019), and, if applicable, incorporating measures of location error, such as dilution-of-precision (DOP) values (Fleming, Drescher-Lehman, et al., 2021). If home range estimation is one of the research targets, the range residency assumption should be verified for all individuals (Fleming et al., 2014a; Silva et al., 2022).

**Fig. 1.**
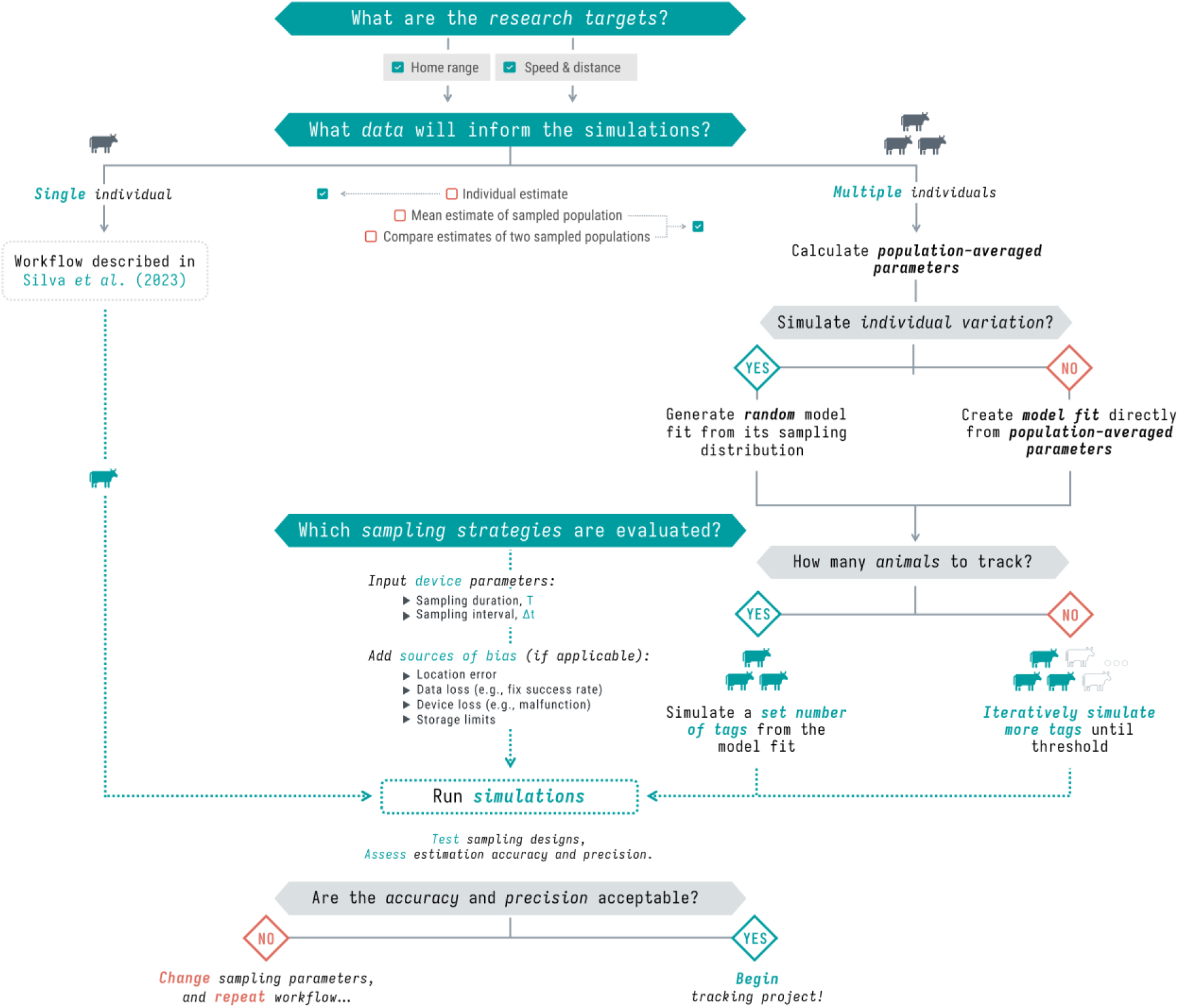
Workflow for evaluating population sample sizes while balancing trade-offs with sampling parameters. The workflow follows a structured approach to evaluate the impact of study design on population-level inferences of key movement ecology metrics. To achieve this, we derive population-averaged parameters from continuous-time movement models fitted to a selected set of individuals. When simulating individual variation, these population-level parameters define a population model from which we generate random model-fit statistics by sampling from the associated sampling distribution. When individual variation is not simulated, we instead create a prototype model with the population-averaged parameters manually specified. This workflow ensures that study designs are tailored to minimize uncertainty and maximize inferential robustness. Beyond the incorporation of population-level inferences, individual-level steps are discussed in further detail in Silva et al. (2023).

Given our assumption of a statistically stationary movement process (*i.e.*, its statistical properties are constant in time over the relevant temporal and spatial scales), it may also be necessary to segment the dataset, by time or behavioral states, prior to parameter extraction (Gupte et al., 2022). Neglecting to address these issues may result in parameter estimates drawn from an unrepresentative sample, undermining the reliability of downstream analyses.

The extraction step can be carried out using standard continuous-time model selection (Calabrese et al., 2016), which evaluates a set of continuous-time stochastic movement processes available within the `‘ctmm’` framework, compares their fit to the data using likelihood-based criteria, and identifies the model and parameter set that best captures the underlying movement patterns. If individual variation is negligible, we can simulate new individuals directly from the extracted parameters. To account for individual variation, however, we simulate new individuals from a mean (uniformly-weighted) model fit estimated using a normal–normal hierarchical framework. This procedure log-transforms scale parameters and applies a matrix logarithm to covariance matrices, followed by a log(χ²) bias correction. It then fits a hierarchical model to estimate population-level means and between-individual variance components, and back-transforms the results with appropriate bias corrections. This approach assumes that the observed distribution of parameters is representative of the true population distribution. However, it explicitly incorporates both within-individual estimation uncertainty and between-individual variability, fully reflecting the uncertainty present in the data. Given that movement behavior may vary considerably within a population (*e.g.*, “behavioral syndromes” or “personalities”; Hertel et al., 2020), the selected dataset should ideally capture a representative range of movement behaviors for the population of interest.

### Running simulations and incorporating sources of bias

After defining clear, quantifiable objectives and extracting all relevant parameters, the next step is to identify which models or sampling schedules we wish to evaluate. This step involves specifying one or more sets of sampling parameters (duration and interval) to be applied in subsequent simulations, conditioned on the movement parameters extracted. During this stage, researchers can generate simulated datasets that account for additional sources of bias and uncertainty that arise during deployment, such as: (1) *location error*, (2) *data loss* (which can reflect the proportion of successful location fixes, *fix-success rate*, or other sources of temporary data loss), (3) *deployment disruption* (which can arise from technical malfunctions, causing a unit to stop recording prematurely and permanently, or from a mortality event), or (4) *storage limits* (GPS units may have finite capacity for storing new locations).

Accounting for location error is of particular importance for speed and distance estimation, especially if the error exceeds the scale of movement between subsequent time steps: as the magnitude of location error, if uncorrelated in time, is independent of Δ*t*, conventional estimation methods (such as straight-line-displacements) will diverge to infinity as sampling frequency increases (Fleming, Drescher-Lehman, et al., 2021; Noonan, Fleming, et al., 2019).

### Estimating individual- and population-level parameters

The next step involves estimating individual-level and population-level metrics of interest. To ensure estimates are comparable while accounting for autocorrelation and other sources of bias (Fleming et al., 2015, 2019; Silva et al., 2022), we use continuous-time methods: home range area is estimated via Autocorrelated Kernel Density Estimation (AKDE; Fleming et al., 2015; Fleming & Calabrese, 2017), while movement speed is estimated through a Continuous-Time Speed and Distance estimator (CTSD; Noonan, Fleming, et al., 2019).

Building upon previous work presented in Silva *et al*. (2023), this workflow now extends beyond a single individual and into analytical targets at the population-level. Specifically, it can evaluate either mean estimates for a single sampled population, or mean estimate ratios for two sampled groups (*e.g.* comparing sexes, ages, habitats). For univariate outputs, such as home range areas and movement speed, individual estimates are treated as having a *χ*^2^ sampling distribution, reflecting the uncertainty inherent in that estimate. The population of estimates is modelled using an inverse-Gaussian (*IG*) distribution (Fleming et al., 2022), which captures both individual-level estimation uncertainty and variation among individuals. For multivariate estimates, a matrix log transformation with a log-χ² bias correction is applied to the individual scale parameters to stabilize variances and correct for small-sample effects. After this transformation, both the sampling distribution and the population distribution are treated as multivariate normal. Finally, model selection is performed on the population variance-covariance parameters to quantify how these parameters vary and correlate among individuals. This modeling framework presents good statistical efficiency even for small *population* sample sizes (*m*), and propagates estimate uncertainty from the individual- to population-level estimates by downweighting uncertain estimates relative to more certain ones (Fleming et al., 2022). From here, we calculate the precision of our estimates for any specified set of sampling parameters and for any number of tracked individuals.

### Performing sensitivity analyses

Beyond assessing estimator bias and precision, researchers must also assess how representative a *simulated* population is relative to the *target* population. While some species may exhibit little inter-individual variation, others may pose significant challenges for study design due to high individual variability (Hertel et al., 2020). Ensuring a robust workflow requires evaluating whether a sample adequately captures this variability, and whether the observed precision of our estimates may arise from the specific individuals sampled (*e.g.*, behavioral outliers, rather than a representative cross-section of the population), particularly at low *population* sample sizes.

To conduct sensitivity analyses, we employ two complementary methods. First, we apply a resampling approach (Adams et al., 1997; Bischl et al., 2012), in which we progressively increase the number of tagged individuals drawn without replacement from the simulated population, starting with two up to the maximum *population* sample size. For each value of *m*, we generate a predefined number of combinations of individuals (*e.g.*, selecting multiple sets of 4 individuals from a total sample of 10) and recompute population-level estimates using the matrix log normal-normal framework. This procedure allows us to assess the stability and precision of our inferences as a function of the inclusion of different individuals at each *population* sample size, and to identify the minimum number of individuals needed to achieve a desired level of precision. Second, we perform a leave-one-out cross-validation approach, where we sequentially exclude each one individual from the simulated population, recalculate the population-level estimate using the matrix log normal-normal framework, and repeat this process until every individual has been left out once. This approach allows us to detect whether any single individual disproportionately affects the robustness of our inferences.

Overall, the entire workflow can be repeated as many times as needed to refine study design, gradually adjusting sampling parameters and/or *population* sample size until the mean estimate error and the uncertainty around it falls within an acceptable threshold. Note that very small *population* samples can limit the reliability of inference, and that any conclusions drawn under such conditions should be interpreted with caution. This threshold is context-dependent and largely determined by the amount of individual variation present in the tracked population.

## Simulations

To further validate the workflow and extend our insights, we performed two detailed simulation studies informed by empirical data: an African Buffalo (*Syncerus caffer*) dataset tracked in Kruger National Park, South Africa, between 2005 and 2006 (Cross et al., 2016), and a Mongolian Gazelle (*Procapra gutturosa*) tracked in the Eastern Steppes of Mongolia between 2007 and 2011 (Fleming, Calabrese et al., 2015). The first set of simulations focused on home range area estimation, while the second focused on movement speed. We ran simulations across progressively larger *population* sample sizes (from two to 50 individuals) for both metrics and both species, systematically varying either sampling duration (for home range estimation) or sampling interval (for movement speed estimation).

To assess our population-level inferences, we conducted both sensitivity analysis described above. For the resampling approach, we randomly drew *m* individuals (ranging from two to 50) for a maximum of 250 combinations of individuals per *m*, and recomputed mean estimates using the matrix log normal-normal modelling framework. To quantify accuracy, we calculated the population mean for all combinations for each *m* and expressed the results as relative errors (%), defined as the difference between the estimated and expected values (under each simulated movement model), divided by the expected value. Negative values indicated underestimation, while positive values indicated overestimation. To quantify precision, we computed the 95% confidence intervals of the mean estimates. We set a ±5% error threshold as a benchmark for reliable inference. For our leave-one-out approach, we also calculated the proportion of estimates that fell within the predefined error threshold of the expected true values (correctness rate). We conducted the simulations on a high-performance computing cluster using `R` version `4.3.3` (R Core Team, 2017), and all subsequent analyses were executed in `R` version `4.5.3` (R Core Team, 2026), with `‘ctmm’` version `1.3.1`, and `‘movedesign’` version `0.3.3`. A step-by-step tutorial of this workflow using the `‘movedesign’ R` Shiny application is available in Supplementary File S2.

## Results

### Home range estimation

For the first simulation study, shorter sampling durations led to an increased underestimation of home range areas for both species, even at large *population* sample sizes. For the African buffalo (*Syncerus caffer*; Fig. 2, top panel), a duration of two months led to an underestimation of 25% at *m* = 50 individuals, with a mean *effective* sample size for home range area of approximately 6.6 (95% CI: 2.7—11.26). This pattern suggests that increasing the *population* sample size does not compensate for insufficient individual-level sampling duration. Achieving an error within the threshold (±5%) for fewer than 50 individuals was only possible for longer durations (≥ 3 years), which yielded lower relative errors and narrower confidence intervals. For a maximum sampling duration of 9 years, we achieved a small overestimation of 1.4% for *N*_*area*_ = 313.5 (195.8—411.9).

**Fig. 2.**
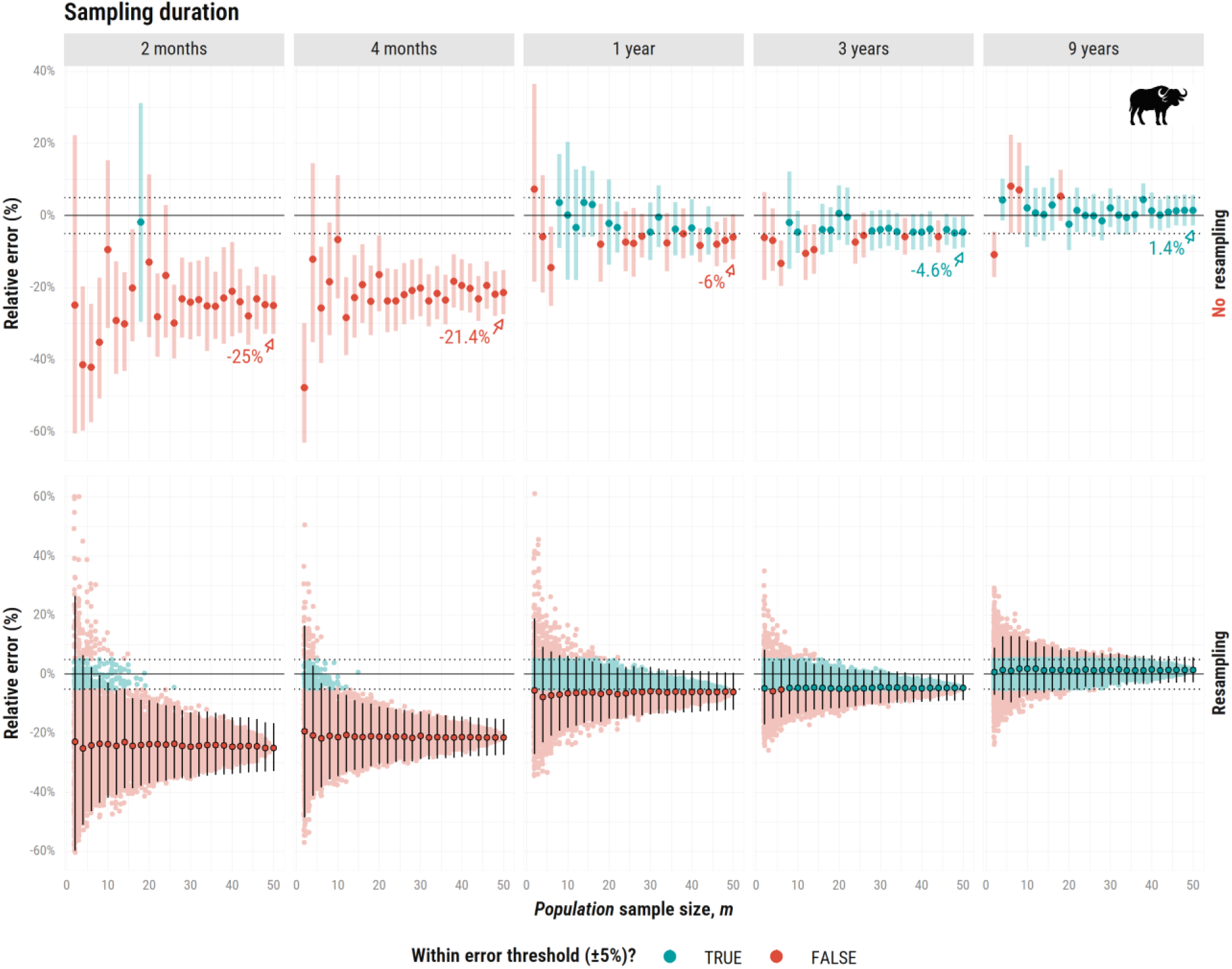
Relative error (%) in home range estimates (AKDE) as a function of population sample size (***m***), ranging from 2 to 50 simulated African buffalos *(Syncerus caffer)*. Each vertical facet corresponds to one of five sampling durations: 2 months, 4 months, 1 year, 3 years, and 9 years. The top panel shows mean estimates based on a single combination of individuals per m (no resampling); estimates in blue are within the error threshold and in red if estimates fall outside of it. The bottom panel shows the resampling approach, where we generated up to 250 random combinations of m individuals, each represented as a colored point; points and vertical lines in black denote the mean point estimates, lower and upper bounds across all combinations. Horizontal dotted lines mark the ±5% error threshold. Very long sampling durations (*e.g.*, 9 years), may be largely impractical or unrealistic, and were used here primarily for demonstration purposes.

Mean estimates varied substantially as *m* increased due to individual variability (Fig. 2, top panel): whether estimates fell within the error threshold sometimes depended on which individuals were sampled, rather than on consistent performance. Resampling stabilized our estimates, and enabled a more reliable assessment of their accuracy and precision (Fig. 2, bottom panel), mitigating the risk of drawing misleading conclusions. At small *population* sample sizes, even the maximum sampling duration resulted in confidence interval widths exceeding the error threshold, underscoring the importance of increasing *m* to improve estimator precision.

In contrast, for the Mongolian Gazelle (*Procapra gutturosa*), underestimation remained severe even at large *population* sample sizes under the same sampling durations as the African buffalo (Fig. 3). While a three-year sampling duration yielded a mean relative error within ±5% for African buffalos, a similarly parameterized scenario led to an underestimation of 9.6% for Mongolian gazelles (*N*_*area*_ = 14.1, 5.5—24.1). Extending to 9 years only marginally reduced error, to an underestimation of 8.1%. Achieving an error within the ±5% threshold is ultimately unfeasible given the median lifespan of Mongolian gazelle (4 years; Olson et al., 2014) and their exceptionally long mean range crossing times of over 2.7 months (1.6—4.5; Fleming, Calabrese et al., 2014). Although Mongolian gazelles are sometimes described as nomadic, their movement is statistically consistent with range residency, albeit at a very large spatial scale.

**Fig. 3.**
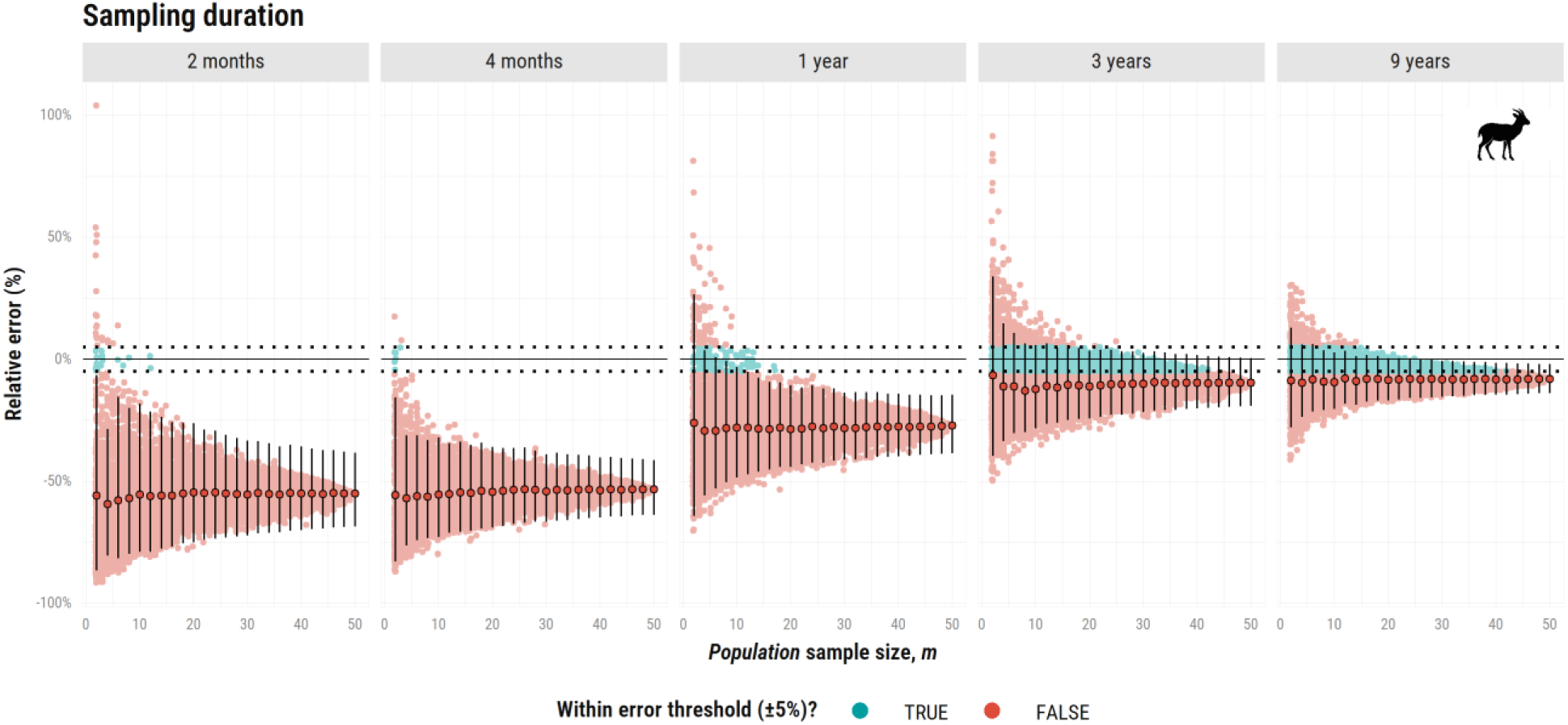
Relative error (%) in home range estimates (AKDE) as a function of population sample size (*m*), ranging from 2 to 50 simulated Mongolian gazelles (*Procapra gutturosa*). This plot presents only the resampling approach (with the mean and confidence intervals across all combinations). The single combination plot is available in Supplementary File S3. Other figure components are consistent with those of the bottom panel of Fig. 2.

Across our leave-one-out iterations (Supplementary File S4), estimates of mean home range area and movement speed showed minimal variation for both species, suggesting that our models were not overly influenced by any single individual. However, only the 9-year sampling duration for the African buffalo yielded a correctness rate of 100% within the predefined error threshold, indicating individual variability no longer impacted our estimates.

### Speed & distance estimation

For the second simulation study, shorter sampling intervals allowed for more reliable movement speed estimates. For the African buffalo (Fig. 4), this scenario yielded a mean relative error within ±1% for the all *population* sample sizes when Δ*t* = 1 ℎ*our* and Δ*t* = 30 *minutes*. For Δ*t* = 30 *minutes*, the *effective* sample size for speed (*N*_*speed*_) averaged 25,621.4 (22,486.0—28,756.7), reinforcing the idea that within-individual sampling effort can compensate for limited *population* sample size when population-level variability is low. At Δ*t* = 4 ℎ*ours*, movement speed was underestimated by 8% even at *m* = 50 (*N*_*speed*_ = 12.0, 8.2—15.7), as we could no longer effectively capture the species’ velocity autocorrelation parameter (*τ*_*v*_ = 32.2 minutes, 22.6— 45.7 minutes).

**Fig. 4.**
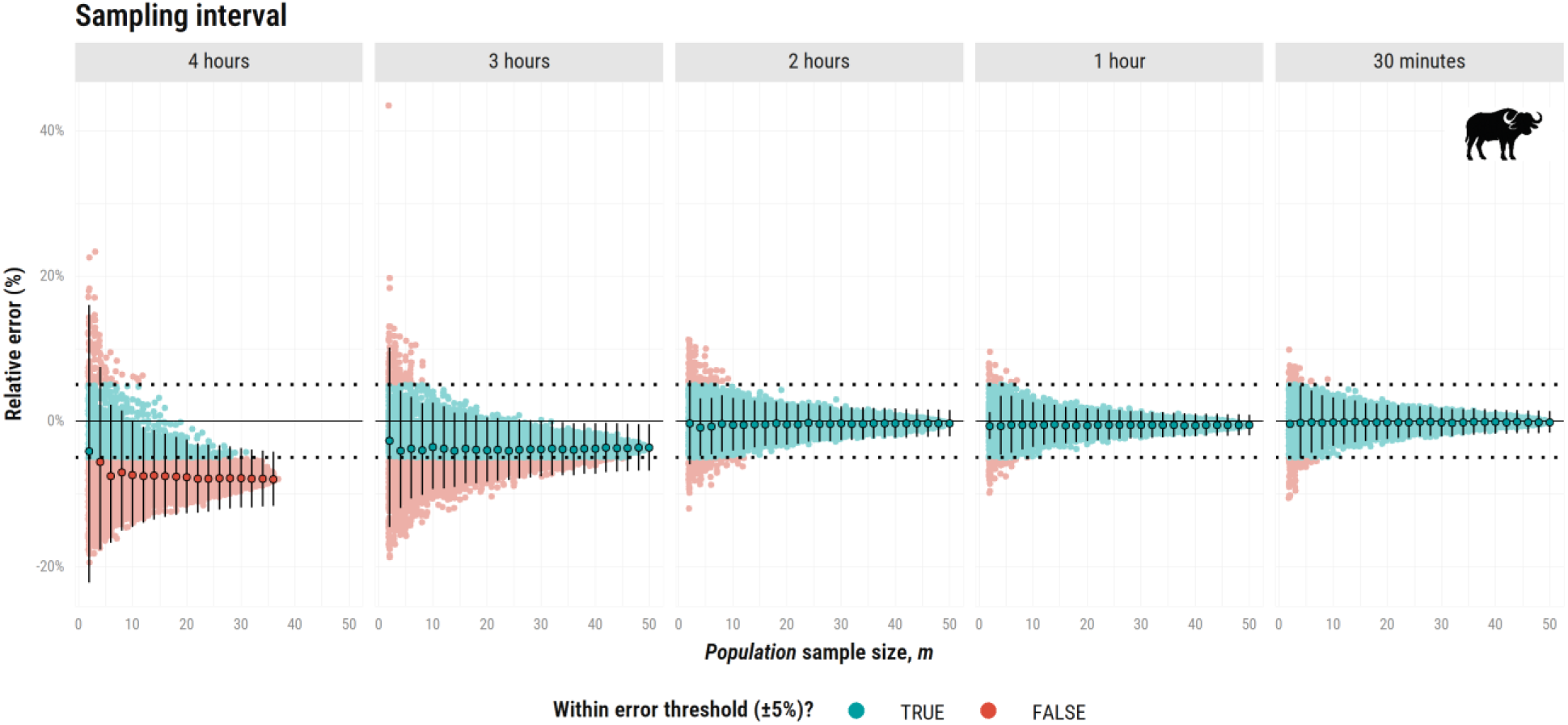
Relative error (%) in speed estimates (CTSD) as a function of population sample size (*m*), ranging from 2 to 50 simulated African buffalos (*Syncerus caffer*). This plot presents only the resampling approach (with the mean and confidence intervals across all combinations). Each vertical facet corresponds to one of five sampling intervals: 4 hours, 3 hours, 2 hours, 1 hour, and 30 minutes. For Δ*t* = 4 hours, estimation was not possible for 13 out of 50 individuals as data was too coarse to support a model with correlated velocity. The single combination plot is available in Supplementary File S3. Other figure components are consistent with those of the bottom panel of Fig. 2.

For the Mongolian gazelles (*Procapra gutturosa*), movement speed estimates also remained highly accurate ranging from long (Δ*t* = 4 ℎ*ours*) to short (Δ*t* = 30 *minutes*) intervals, with a mean relative error always within the ±1% threshold (Fig. 5). This consistency stems from the species’ velocity autocorrelation parameters (*τ*_*v*_ = 0.8 days, 0.4—1.3 days), which enabled reliable speed estimation even at relatively coarser resolutions, or smaller *population* sample sizes. Ultimately, for Mongolian gazelles, further decreasing sampling interval yielded minimal gains for speed estimation, but could significantly compromise home range estimation given the trade-off between battery life and resolution of some biologging devices (Silva et al., 2023).

**Fig. 5.**
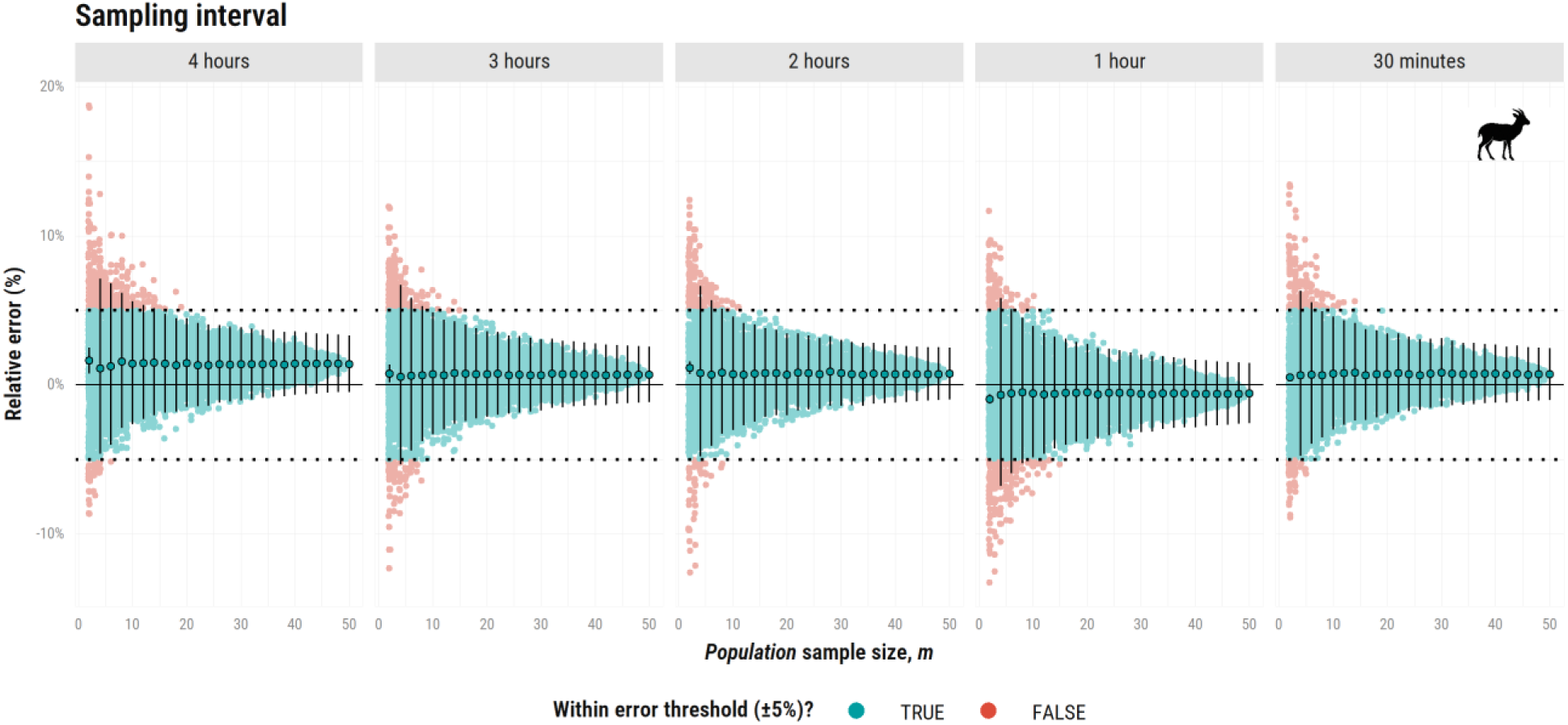
Relative error (%) in speed estimates (CTSD) as a function of *population* sample size (***m***), ranging from 2 to 50 simulated Mongolian Gazelles (*Procapra gutturosa*). This plot presents only the resampling approach (with the mean and confidence intervals across all combinations). The single combination plot is available in Supplementary File S3. Other figure components are consistent with those of the bottom panel of Fig. 4.

For our leave-one-out approach (Supplementary File S4), both species showed highly consistent performances. The Mongolian gazelle achieved a correctness rate of 100% across all sampling intervals, while the African buffalo only failed for the 4-hour interval (correctness rate of 0%), once again highlighting the mismatch between this sampling resolution and the species’ underlying velocity autocorrelation timescale (Δ*t* ≫ *τ*_*v*_).

## Discussion

Tracking technology can offer valuable insights into fine-scale and large-scale processes of animal movement behavior and space-use requirements, yet researchers face numerous challenges during study design and deployment. The value of a research project to managers and conservation practitioners is partially determined at its inception, when reliable data-driven knowledge shapes study design. This crucial step ensures that the project has the best possible chance to yield meaningful results that will correctly inform future conservation strategies or management interventions (Rüegg et al., 2014; Urbano et al., 2024). Considering the substantial time and financial investment required for any animal tracking project, researchers should carefully consider study design and adjust as needed to manage error, limit impacts on animal welfare, and reduce the risk of misinterpreting ecological patterns. This includes selecting tracking devices that comply with widely accepted guidelines, such as the 5% body weight rule, or reducing deployment duration to mitigate potential physiological or behavioral impacts on the animals. This is especially critical for threatened species, where biases can have serious consequences for conservation efforts and long-term population viability (Noonan et al., 2020). For instance, in the context of human-wildlife conflict, overestimating home range areas could result in the misclassification of certain animals as “problematic” (Bauer & De Iongh, 2005). Meanwhile, underestimating home range areas could inflate density estimates (Rinehart et al., 2014), or overlook potential anthropogenic impacts (Benitez et al., 2022).

Reliable estimates must be both accurate and precise. While larger *population* sample sizes improved estimate precision, individual-level sampling parameters were critical for accurately estimating mean home range or speed (*e.g.*, longer durations or shorter intervals, respectively). However, very long sampling durations (*e.g.*, *T* = 9 *years* as presented in our simulations), may be largely impractical or unrealistic, and were used here primarily for demonstration purposes. As sampling duration increases, so does the risk of device failure or loss of tracked individuals. For study designs that rely on long sampling durations for reliable estimates, it is crucial to account for these potential failures and integrate them into the simulations. The `‘movedesign’` application allows users to quantify these and other limitations, providing a more comprehensive evaluation of long-term tracking studies. When the risk of device failure or animal mortality during deployment is high, we recommend proportionally increasing population sample sizes to compensate for these losses.

Researchers also strive for sampling strategies that address multiple research questions. However, achieving an optimal balance between competing goals can be challenging. As demonstrated in our case study (Supplementary File S1), a single approach may offer a compromise, yet remain only marginally effective in reliably estimating both home range and movement speed. Rather than refining one aspect of the project, such an approach risks returning unreliable results for both quantities. In many cases, no universally optimal sampling strategy exists, particularly when the position and velocity autocorrelation timescales diverge substantially (Noonan et al., 2019). Because home range crossing time typically spans days to months, while directional persistence spans minutes to hours (Noonan et al., 2019), a given sampling frequency may be too coarse to resolve fine-scale movement patterns, or the sampling duration too short to yield reliable estimates of home range areas.

A similar issue arises when using movement data to define protected areas, which may stem from neglecting autocorrelation (Noonan, Tucker, et al., 2019), or from improperly designed studies that result in very low effective sample sizes. In addition, incorrectly detecting differences between two tracked groups when none exist, or failing to detect differences when they do, could also reduce the effectiveness of management efforts (particularly when comparing individuals within protected *vs.* human-dominated landscapes).

Optimal study design is target-dependent, as reducing estimate bias relies on how well sampling parameters capture the movement behavior of the *target* population, which is strongly governed by the relationship between the sampling parameters and the sampled individuals’ autocorrelation timescales. Both the frequency of behavioral state transitions and the influence of external environmental drivers must also be considered (Hertel et al., 2020), as both play significant roles in shaping animal behavior. Researchers must carefully select datasets that accurately represent the target population, and the optimal study design should not compromise estimate precision to ensure informative outputs (Nakagawa et al., 2024). Dealing with small effective sample sizes requires caution, and this risk should be minimized, even when applying bias mitigation measures (Fleming et al., 2019; Silva et al., 2022). However, significantly increasing sample size may be neither feasible nor ethical. For Mongolian gazelles, which have a home range crossing time spanning months, obtaining a sufficiently large effective sample size is impossible, regardless of study duration, due to their short lifespan. Alternatively, achieving a sufficiently long sampling duration may require tag weights that exceed the recommended 5% of the animal’s body mass for some species (Kenward, 2000). Such an approach raises considerable ethical concerns and should be avoided to uphold best practices in animal welfare.

Tracking a representative sample of individuals presents another stumbling block, but once again no universal solution exists. Our workflow allows researchers to systematically assess any sampling constraints on a case-by-case basis, offering valuable insights into estimator precision and supporting informed decision-making. Movement ecology, akin to other fields with practical applications, must maximize information acquisition for conservation (Milner-Gulland & Shea, 2017). Our findings underscore the critical interaction between species parameters and sampling parameters in determining the optimal *population* sample size. The bias introduced by shorter sampling durations or larger sampling intervals outweighed the bias associated with tracking fewer individuals for the estimation of both home range and speed. Shorter sampling durations may not provide reliable estimates of mean home range areas, regardless of the *population* sample size. In contrast, prioritizing shorter sampling intervals ensures sufficient data resolution for accurate speed estimation even at low *population* sample sizes. Researchers studying species with shorter home range crossing times have more flexibility in choosing sampling durations, whereas those studying species with longer directional persistence timescales have greater flexibility in selecting sampling intervals. In general, we recommend prioritizing biologgers and sampling strategies that reduce estimation error, as this approach is often more effective than simply increasing the number of individuals tracked. Examples include longer-lasting batteries to allow longer sampling durations for home range estimation, or higher frequency sampling for speed estimation. When research goals focus on comparisons across groups, or on populations with high individual variation, ensuring adequate representation may justify allocating resources to track additional individuals.

We also observed that for very small *population* sample sizes (*m* < 4 individuals), individual variation causes point estimates to be highly variable and can sometimes produce deceptively low relative errors, even when the average estimate is far from the true value. Any conclusions drawn from such small sample sizes are unlikely to generalize well to the broader population, regardless of the methods used. Ultimately, we cannot discount the importance of individual variation in movement behaviors (Hertel et al., 2021). Factors such as age, sex, or reproductive status can all contribute to this variation, resulting in distinct space-use strategies among individuals.

The workflow presented here is built on the Ornstein-Uhlenbeck with Foraging (OUF) model (Fleming et al., 2014a, 2014b), which features correlated positions and velocities. This OUF model is currently the most frequently selected across empirical GPS datasets (Fagan et al., 2025; Noonan et al., 2019). However, as a result, all inferences are conditional on assumptions of stationarity and, for home range estimation, range residency. Apparent non-stationarity often reflects transitions between distinct movement behaviors, such as dispersal or migration. When such behaviors are identified, users should segment the data and evaluate each resident period separately to ensure that the model assumptions are satisfied. When individuals exhibit fundamentally different movement regimes (*e.g.*, a population with both migratory and range resident animals), model-based inference pertains only to the subset satisfying the model assumptions. In such cases, estimates should be interpreted as parameters of the resident subpopulation rather than the entire population. In addition, the continuous-time movement model framework is modular, allowing additional movement processes to be incorporated into the workflow as they become available, thereby facilitating adaptation to new movement behaviors, or ecological contexts. However, the recommendations presented here are currently specific to home range and movement speed estimation, and should not be extended to other metrics without further validation.

The allocation of finite resources must be weighed against each additional tracked individual, as larger *population* sample size may lead to diminishing returns. The key trade-off in study design lies between optimizing within-individual data quality and ensuring adequate representation of the individual variation present in the target population. For both very large *effective* and *population* sample sizes, population-mean estimates will approach the expected true values even with conventional methods (such as unweighted sample-mean analysis or Kernel Density Estimators; Fleming et al., 2022; Silva et al., 2022); however, even under these conditions, it is better to employ sampling-insensitive methods that adhere to the assumptions of modern movement data, report uncertainty, and can be easily evaluated.

A one-size-fits-all approach to movement ecology study design is insufficient; instead, a flexible framework that incorporates data-informed simulations yields the most actionable insights. When designing a tracking study, some prior knowledge of the species’ movement is necessary to make informed decisions. If representative pilot data are unavailable, users can use rough estimates based on expert knowledge, similar species, or plausible ranges of movement parameters to simulate likely outcomes. If pilot data are available, these can be incorporated directly to inform study design. While we recognize that these approaches provide only rough estimates, they still allow researchers to evaluate and refine study designs rather than moving forward without any guidance. The availability of movement data is rapidly increasing (Kays et al., 2022), providing researchers with even more opportunities to inform study design even when species-specific pilot data are limited.

By creating and sharing this workflow, we aim to establish a feedback loop that enhances the integration of movement ecology data into conservation efforts, refining study design and improving the reliability of animal tracking projects. Crucially, it can support researchers in preparing grant proposals and permit applications, providing clearer insight into the sampling effort required to funding agencies and regulatory bodies. And as the continuous time movement model framework underlying this workflow expands, it will naturally accommodate additional analyses over time.

## Data Availability statement

The empirical datasets used in the manuscript are openly accessible: the African buffalo and Mongolian gazelle tracking datasets are included in the `‘ctmm’ R` package (Fleming & Calabrese, 2022). Source code of the `‘movedesign’ R` package and Shiny application is available on `GitHub` (https://github.com/ecoisilva/movedesign), and version `0.3.3` has been archived on `Zenodo`. All generated datasets and `R` scripts required to reproduce the analyses and figures in this manuscript will be made available on `GitHub` (https://github.com/ecoisilva/studydesign_ms).

## Acknowledgements

This work was partially funded by the Center of Advanced Systems Understanding (CASUS), which is financed by Germany’s Federal Ministry of Education and Research (BMBF) and by the Saxon Ministry for Science, Culture and Tourism (SMWK) with tax funds on the basis of the budget approved by the Saxon State Parliament. C.H.F., J.M.C., and W.F.F. were supported by NSF IIBR 1915347.

## Authors’ contributions

I.S., C.H.F., and J.M.C. led conceptualization; I.S. led simulations, analyses, and visualization; I.S. and J.M.C. led the writing of the manuscript; I.S., C.H.F., M.J.N., W.F.F. and J.M.C. edited the draft. All authors contributed critically to the drafts and gave final approval for publication.

