## Supplementary material for "Too few, too many, or just right? Optimizing sample sizes for population-level inferences in animal tracking projects": Suplementary File S1

### **S1.** Case study

This document describes a case study using the workflow presented in the main manuscript. Our goal was to obtain a reliable estimate (within a 5% error threshold) of both **mean home range area** and **mean movement speed** of a tagged population of African buffalos (*Syncerus caffer*) in a protected area. We wished to limit expenses by acquiring the minimum number of tags needed to obtain reliable mean estimates for the tagged population, while accounting for individual variation.

To inform our analyses, we used a GPS movement dataset of African buffalos tracked between 2005 and 2006 in Kruger National Park, South Africa (Cross et al., 2016), available on Movebank and with the **'ctmm'** package. We assessed range residency by examining the variograms of all six individuals within this dataset. Variograms allow for a cursory confirmation of range residency by checking if the semivariance function reaches an asymptote (Calabrese et al., 2016; Fleming, Calabrese, et al., 2014; Silva et al., 2022). All but one (named "Gabs") fit the range residency assumption, so we extracted population-level species parameters using the remaining five individuals.

When choosing a suitable tag for our focal species, we adhered to the standard guideline that units should not exceed 5% of an animal's body mass. With an average African Buffalo weighing 567.5 kg, weight constraints are negligible as most commercially available GPS collars are well below the 5% body mass threshold. Given a set budget of $20,000 USD, we evaluated three competing sampling schedules, with the goals of estimating both mean home range area and movement speed:

— **Option 1**: Sampling duration of **1 year**,
with a new location collected **every 15 minutes** (≈ 96 locations/day);
**Model A**, high cost (≈ $2500 USD), tracking a maximum of 8 individuals.

— **Option 2**: Sampling duration of **6 years**,
with a new location collected **every two hours** (≈ 12 locations/day);
**Model A**, high cost (≈ $2500 USD), tracking a maximum of 8 individuals.

— **Option 3**: Sampling duration of **4 years**,
with a new location collected **every day**;
**Model B**, low cost (≤ $1000 USD), tracking a maximum of 20 individuals.

The relative error in population-level mean home range area and mean speed & distance estimates varied across the three competing sampling schedules (**Figure S1.1**). For home range estimation, **options 2** and **3** achieved a mean error within the ±5% threshold, with underestimations of 3.3% and 3.1%, respectively. For speed and distance estimation, **options 1** and **2** met this criterion, with their 95% confidence intervals also falling within the set threshold.

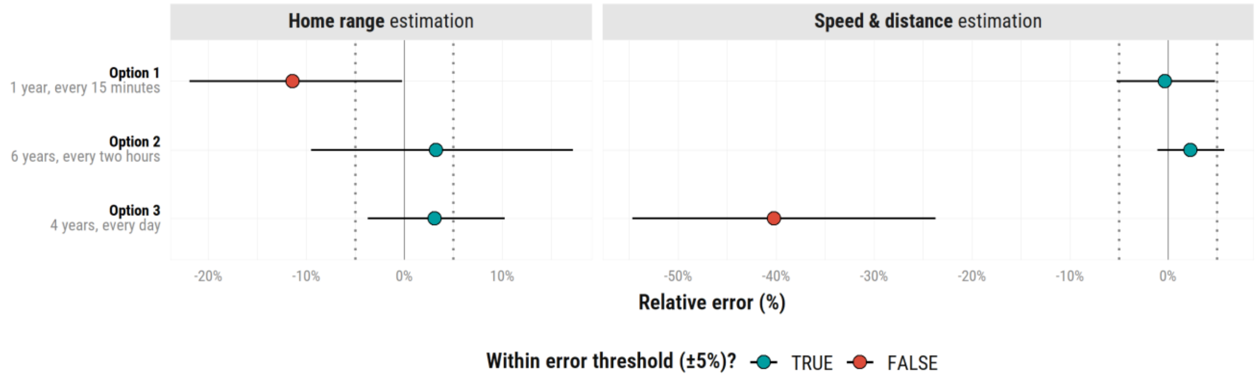


**Figure S1.1**. **Study design comparisons for a simulated population of African buffalos (*Syncerus caffer*).** Relative error (%) in home range estimation (left) and speed & distance estimation (right) for three sampling schedules**.** Colored points and horizontal bars represent mean error ±95% confidence intervals, with values falling within the error threshold in blue, and those outside of it in red. Vertical dotted lines mark the ±5% error threshold.

Finally, we conducted sensitivity analyses for each of our sampling schedule options. With the resampling approach for home range estimation (**Figure S1.2**, top panel), we can see that reducing the number of deployed tags further is ill-advised, as high individual variation means that lower *population* sample sizes have high uncertainty. For example,

**options 2** and **3** resulted in relative errors within the ±5% threshold, with underestimations of 3.3% and 3.1%, respectively. For speed and distance estimation (**Figure S1.2**, bottom panel), **option 3** failed to produce reliable results, underestimating speed by over 43%. It should be noted that estimation was not possible for 17 out of 20 individuals as the data was too coarse to support a model with correlated velocity (Noonan et al., 2019). Both **options 1** and **2** remained within the ±5% error threshold, with only a marginal difference between them, with **option 2** achieving the lowest relative error (0.1%) at $m = 8$.

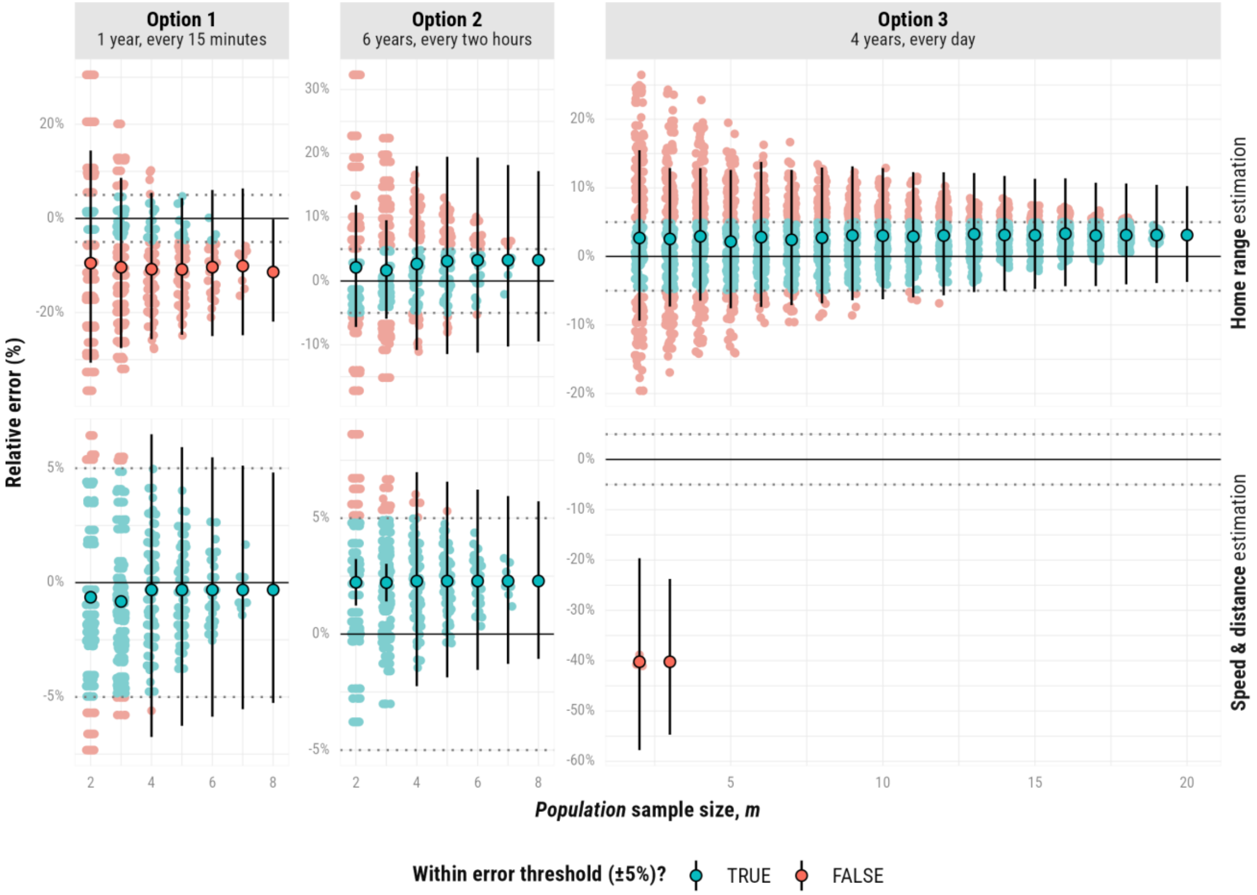


**Figure S1.2. Resampling approach to assess the minimum number of tagged African buffalos (*Syncerus caffer*) needed for reliable estimates.** Panels show relative error (%) in home range (AKDE, top) and speed & distance estimates (CTSD, bottom) across three sampling strategies as a function of *population* sample size ($m$). Horizontal dotted lines mark the ±5% error threshold. We generated 250 random combinations of $m$ individuals, each represented as a colored point: in blue for estimates within the error threshold and red for estimates outside of it. Points and vertical lines in black denote the mean and confidence intervals across combinations.

With our leave-one-out approach (**Figure S1.3**), and for home range estimation, **option 2** exhibited greater variability in the estimate error (mean = 3.8%, SD = 2.9%) than **option 3** (mean = 3.1%, SD = 0.8%). Furthermore, **option 2** had a lower correctness rate (62.5%) compared to **option 3**. These findings highlighted that **option 2** was more sensitive to each new individual included in the meta-analysis.

Only one of our sampling schedules (**option 2**: 6-year duration, 2-hour interval, and 8 individuals) satisfied the specified error threshold for both research targets. Nevertheless, wide confidence intervals and low correctness rate suggested that a higher *population* sample size would yield more reliable estimates. These finding suggested that it may be necessary to prioritize one research target over the other, or consider alternative sampling schedules.

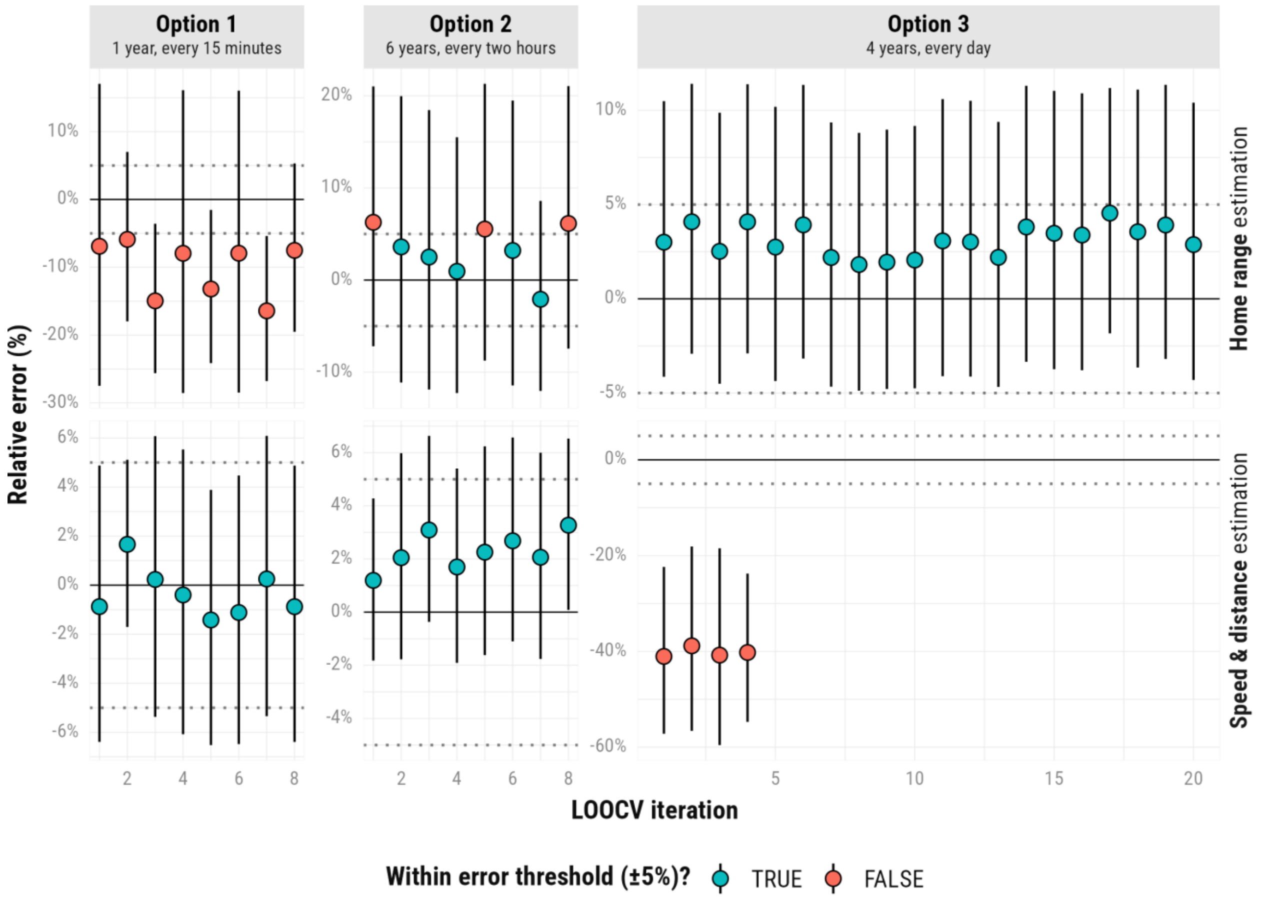


**Figure S1.3. Leave-one-out approach for three competing sampling schedules.** Mean relative errors (colored points) and 95% confidence intervals (bars) are shown, with values falling within the ±5% error threshold highlighted in blue, and those outside of it in red.

The decision to prioritize home range or speed/distance estimation should depend on the ecological context. For example, if the goal is to determine long-term space-use requirements (such as for the delineation of protected areas or the design of management interventions) ***home range estimation*** is preferable. However, for some species, fully resolving home ranges may require tracking durations that exceed typical grant timelines, making it logistically challenging. Conversely, if the focus is on understanding short-term movement responses to factors such as land-use changes, human activity, or linear infrastructures, then estimating ***speed and fine-scale movement dynamics*** is more appropriate. Ultimately, users should balance these considerations to design a study that most effectively informs conservation, management decisions, or research priorities for the species of interest.
