## Supplementary material for "Too few, too many, or just right? Optimizing sample sizes for population-level inferences in animal tracking projects": Suplementary File S2

### S2. Tutorial for `‘movedesign’`

This document describes in detail how to run the workflow presented in main manuscript using the `‘movedesign’` `R Shiny` application. We will outline the process by which we assess the impact of different sampling designs on common movement ecology metrics, *i.e.*, home range area and movement speed, for population-level inferences. All code has been executed in `R` version `4.5.3` (R Core Team, 2026), `‘ctmm’` version `1.3.1`, and `‘movedesign’` version `0.3.3` (Silva, 2025).

To begin, open `R` or `RStudio`, and run the following lines of code:

```
# Install the package first:
install.packages("movedesign")

# Load the package and start the Shiny application:
library(movedesign)
movedesign::run_app() # will open the app
```

To illustrate the workflow, we will apply it to simplified case studies, designed for quick execution. We will use a GPS tracking dataset of African buffalos (*Syncerus caffer*) tracked in Kruger National Park between 2005 and 2006 (Cross *et al.*, 2016), to inform our simulations. Our primary goal is to reliably **estimate mean home range area and** mean **movement speed** of a population of African Buffalos.

We begin by selecting the appropriate workflow in the `‘Home’` tab. In this tutorial, we will showcase two workflows: one for **deploying a set number of units**, and another for determining the **minimum number of units** required to achieve our targets. First, we specify our **data source** (in this case `‘Select’`, as this dataset is included within the `‘ctmm’` package). For both workflows, we tick the `‘☐ Add individual variation’` checkbox. This allows us to account for individual differences, rather than assuming all individuals behave identically.

### **S2.1.** Deploying a set number of units:

For this workflow, we select both **research targets** (home range, and speed & distance estimation), and **analytical targets** (we wish to compare estimates of two sampled groups), as well as the other options mentioned above.

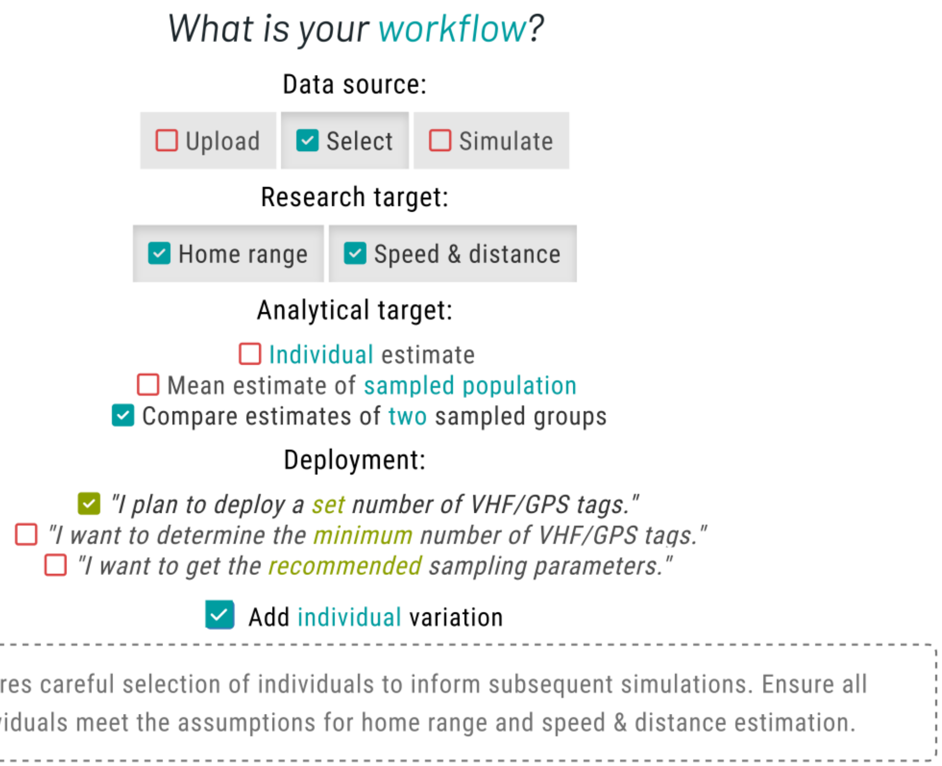


Next, we navigate to the **`'Select data'`** tab (left sidebar), where we can access the movement datasets available within the application. Here, we choose the **African buffalo** dataset, which includes tracking data for six individuals: **`Cilla`**, **`Gabs`**, **`Mvubu`**, **`Pepper`**, **`Queen`**, and **`Toni`**.

To inform the initial simulations, we can extract parameters from a single individual, or from multiple individuals. For a single individual, you can follow along the workflow described in Silva *et al.* (2023). However, it is now possible to extract population-level species parameters ($\tau_p$, $\tau_v$, and $\sigma_p$) from multiple individuals, so we select all African buffalos from the dropdown menu[1].

Before proceeding further, you should visually inspect the variograms to confirm range-residency through the **`'Variogram'`** tab of the **`'Data Visualization'`** box. Variograms allow users to check if

[1] Users may also select individuals directly from the table or the plot in the **`'Data Visualization'`** box.

semivariance reaches an asymptote, and facilitate a cursory confirmation of range residency (Calabrese *et al.*, 2016; Silva *et al.*, 2022). Downstream results may be unreliable if we violate the range residency assumption as, from this point onward, the ‘`movedesign`’ application will operate under the assumption that the data originates from a range-resident species.

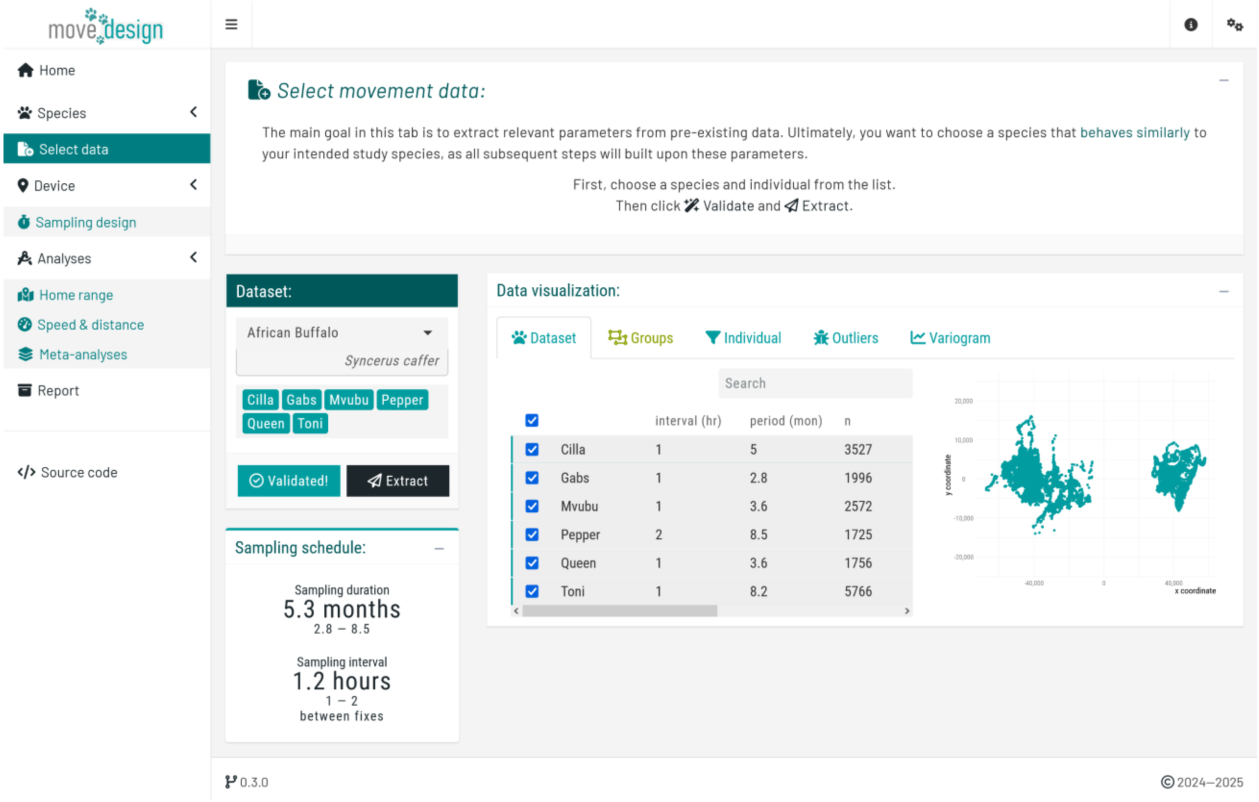


For this tutorial, we will proceed with all six individuals. In a real workflow, however, we would exclude `Gabs` and proceed with the remaining individuals. Next, we validate the selection by clicking the ‘`Validate`’ button (which needs to switch to ‘`Validated!`’) and, afterwards, the ‘`Extract`’ button to retrieve species parameters. Upon successful extraction, the ‘`Displaying parameters`’ box presents our extracted species parameters: the **position autocorrelation** ($\tau_p$) and the **velocity autocorrelation timescale** ($\tau_v$). For the African buffalos, the mean $\tau_p$ is `10.1 days` (`95%` CI: `6.9`, `14.7`), and a mean $\tau_v$ is `32.5 minutes` (`95%` CI: `24.9`, `42.6`). These parameters serve as the foundation for all subsequent simulations as we evaluate study design.

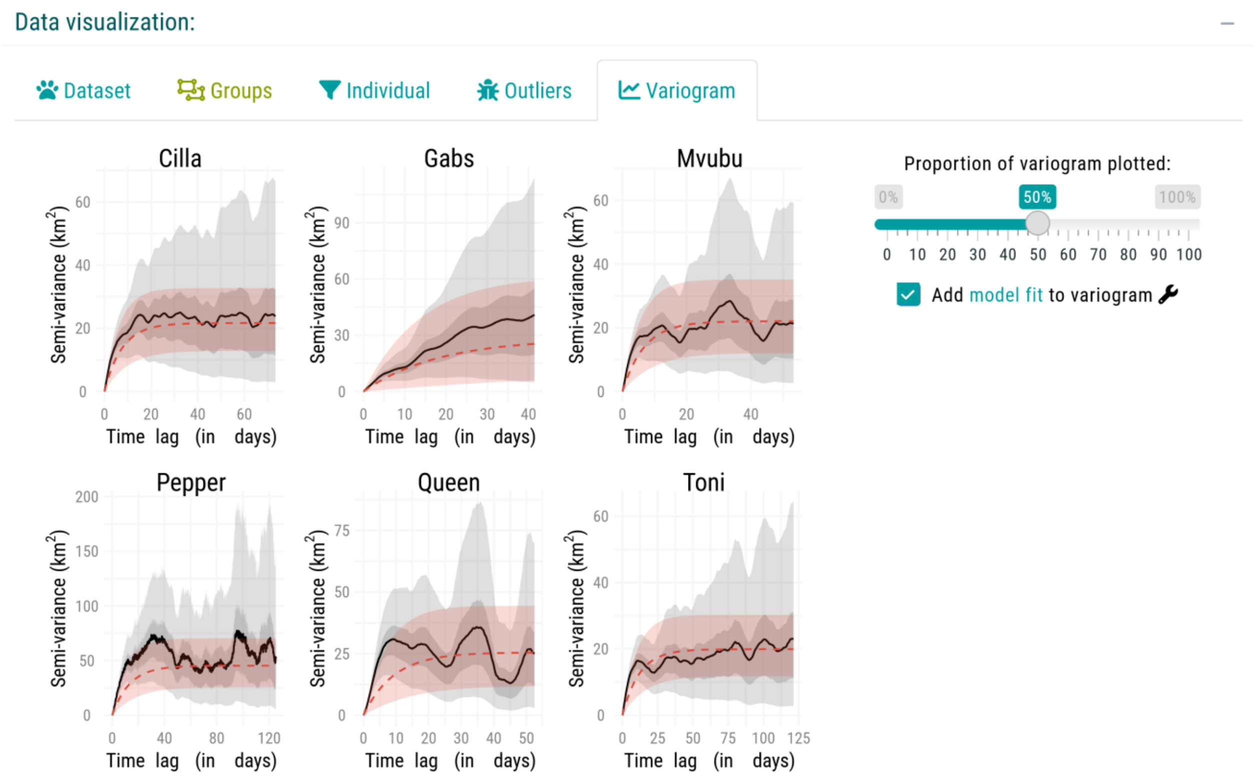


Next, we need to select the groups that will guide both sets of simulations. For this workflow, we will use dummy groups (placeholders with no biological significance), to demonstrate how you can apply it to your own case study. Go to the **'`Groups`'** tab, also available in the **'`Data Visualization`'** box:

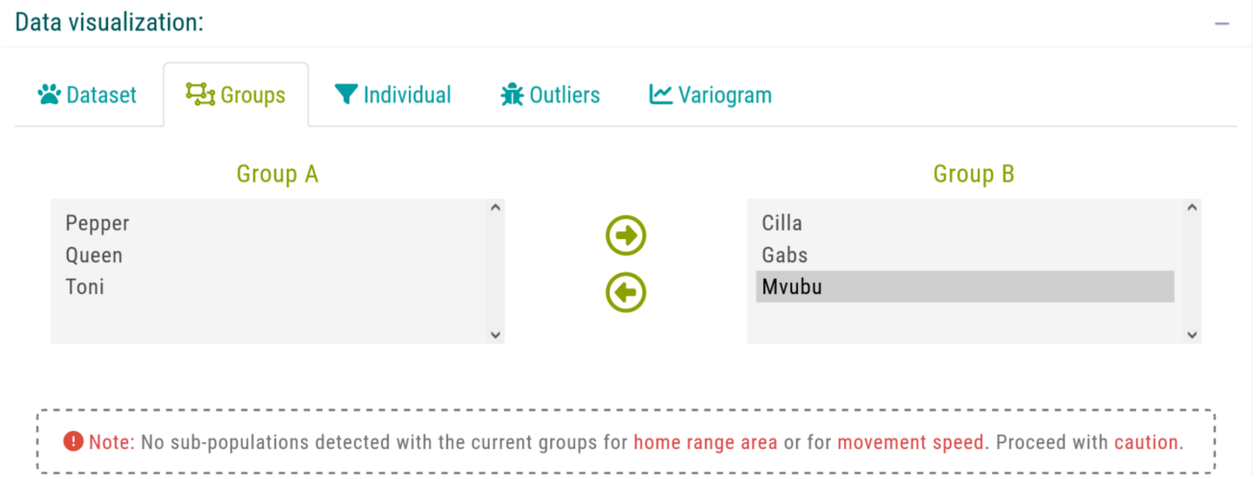


We assign `Pepper`, `Queen`, and `Toni` to **Group A**, and `Cilla`, `Gabs`, `Mvubu` to **Group B** (double-clicking on an individual automatically moves it to the other group). Parameters will then be extracted separately from each group, to inform group-level estimates.

Next, we navigate to the `‘Sampling design’` tab, where we input our sampling parameters. For this tutorial, we will consider the following sampling schedule: a ***sampling duration*** of `3 months`, with 12 new locations collected per day (***sampling interval*** of `2 hours`). To set this particular schedule, we ensure that `‘GPS/Satellite logger’` is selected in the first dropdown menu, and untick the `‘☐ Select from plot’` checkbox in the `‘Device settings’` box before proceeding, which will prompt us to manually input the sampling interval. Then, we set `‘GPS battery life’` (equivalent to the maximum **sampling duration**) to `3 months` and `‘What sampling interval will you evaluate?’` to `2 hours`.

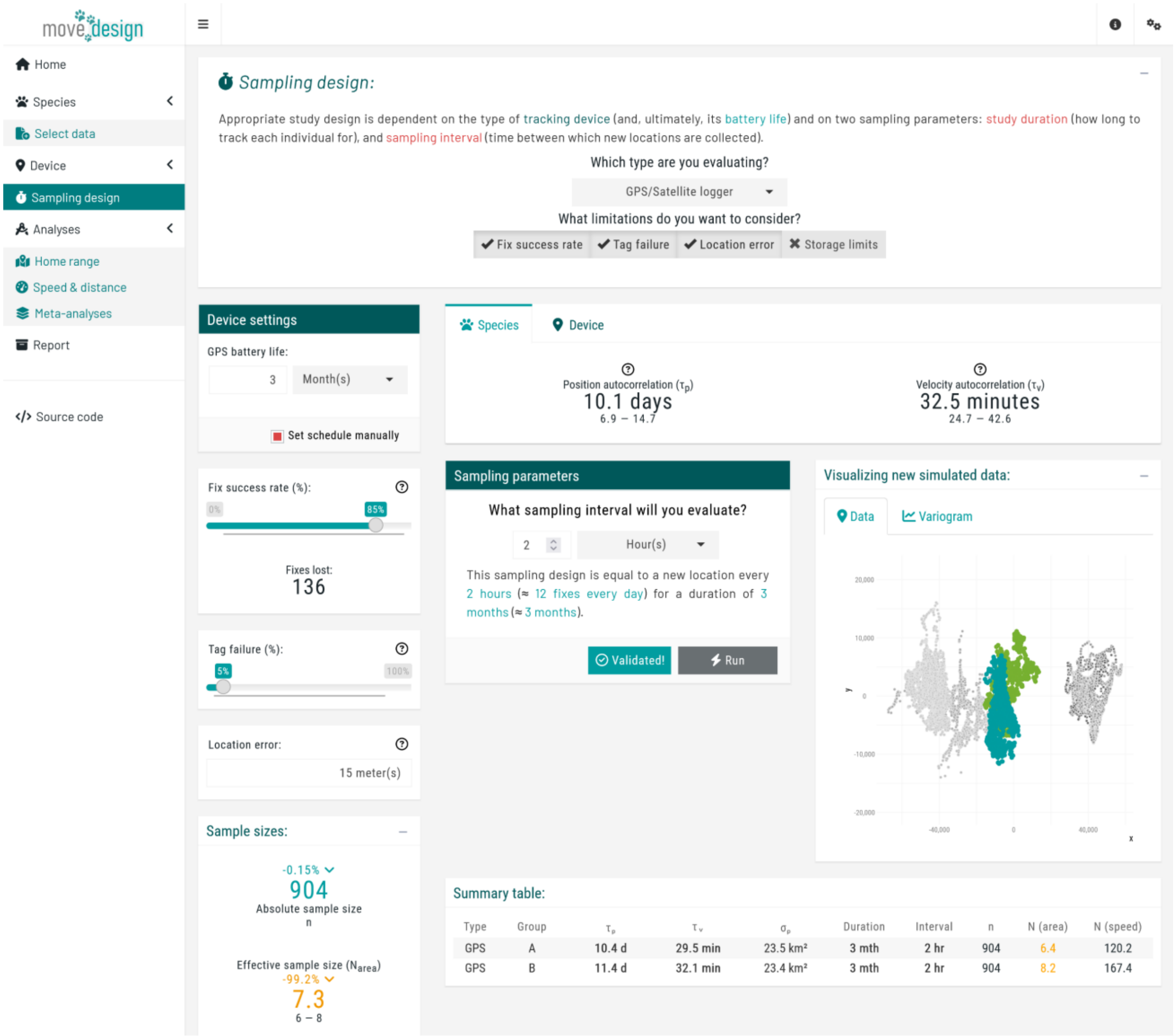


We incorporate three additional components: **fix success rate** (here expected to be, on average, `85%`), reflecting the reliability of the GPS signal; **tag failure** (`5%` chance of a tag failing during data collection); and **location error** (averaging `15 meters`). At this point, we click the `‘Validate’` button (verifying once again that it switches to **`'Validated!'`**) and, afterwards, the **`'Run'`** button. Once a message appears confirming that this step has been successfully completed, we proceed to the **`'Home range'`** tab below **`'Analyses'`**.

To start the estimation process, we click the **`'Run estimation'`** button located in the top box. We can now see the outputs for a single simulation, providing a starting point. The relative error in home range area is an underestimation of **46%**. Note that due to the randomized nature of seed generation and the inclusion of individual variation, values may differ substantially across runs.

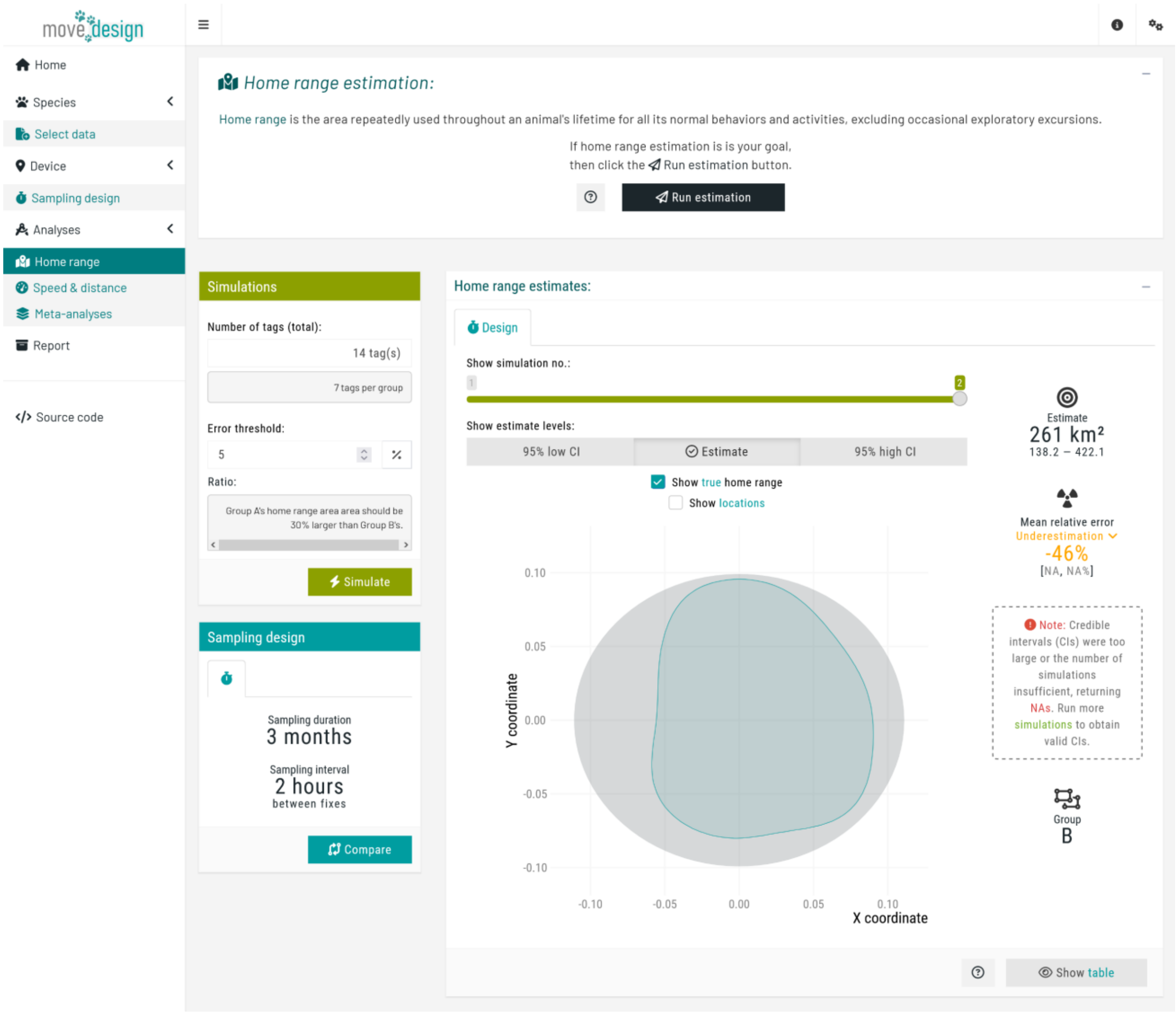


During this workflow, as we wish to compare estimates of two sampled groups, the application always simulates two individuals at a time: one from **group A**, and another from **group B**. We can view the outputs for each simulation using the **`'Show simulation no.:'`** slider. Most importantly, in the **`'Simulations'`** box (top right corner), we set the **total number of tags** and the **error threshold**.

This error threshold serves as a reference point in subsequent plots. It enables users to determine whether a given study design remains within the acceptable error range. We set this threshold to 5%, so any design that ultimately produces an estimate with a relative error above 5% exceeds the predefined tolerance and requires adjustment. We then set the **total number of tags** to 14 individuals, before clicking the 'Simulate' button. A message then appears, indicating the expected runtime; we must wait for this process to complete before we can explore all the outputs.

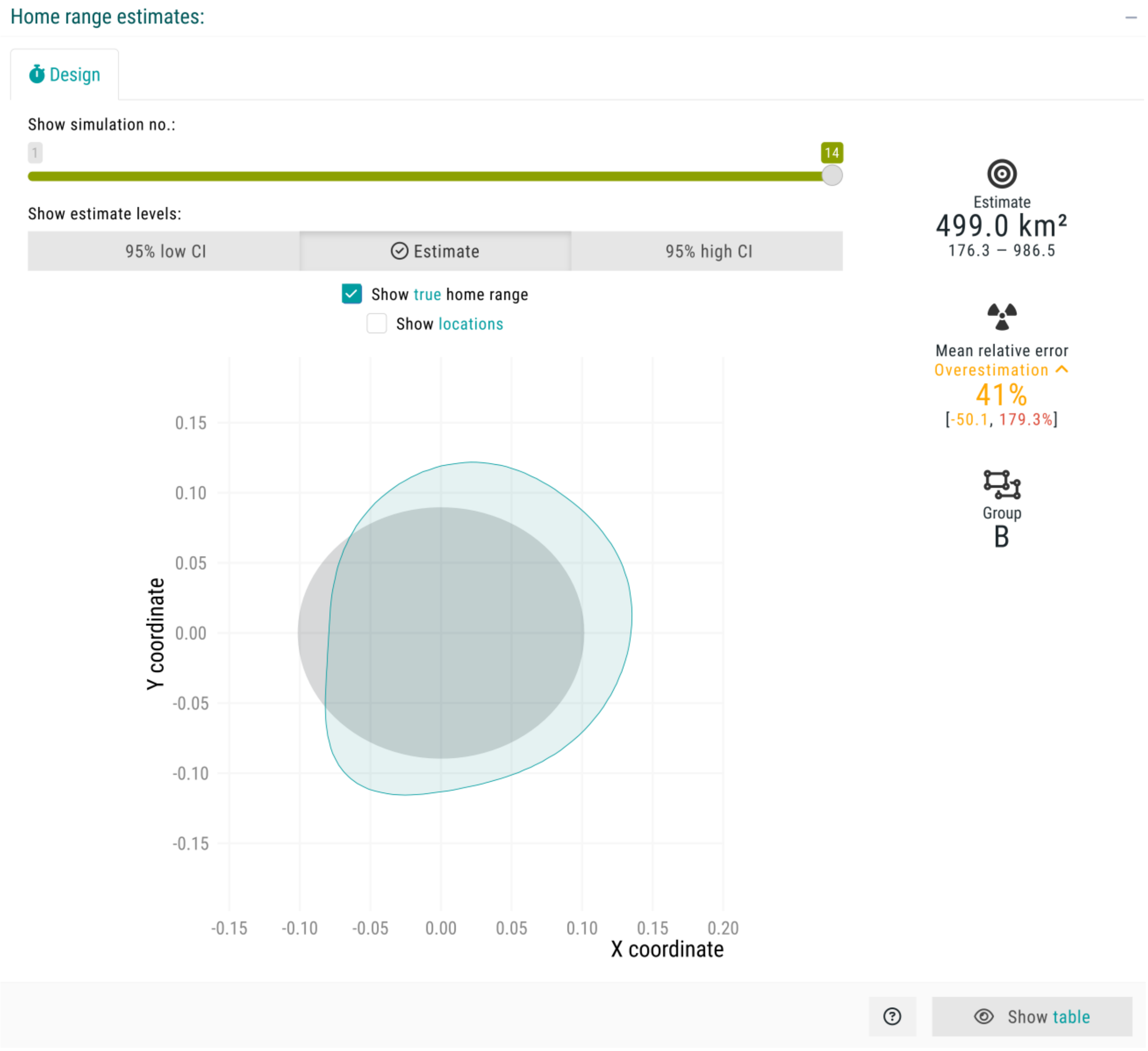


Then, we proceed to the 'Speed & distance' tab, clicking the 'Run estimation' button to obtain estimates of **speed & distance traveled**. Overall, the **individual-level home range areas** across all 14 individuals were underestimated by 17% (no CIs due to low sample size), while the **individual-level movement speeds** were overestimated by 3.9%.

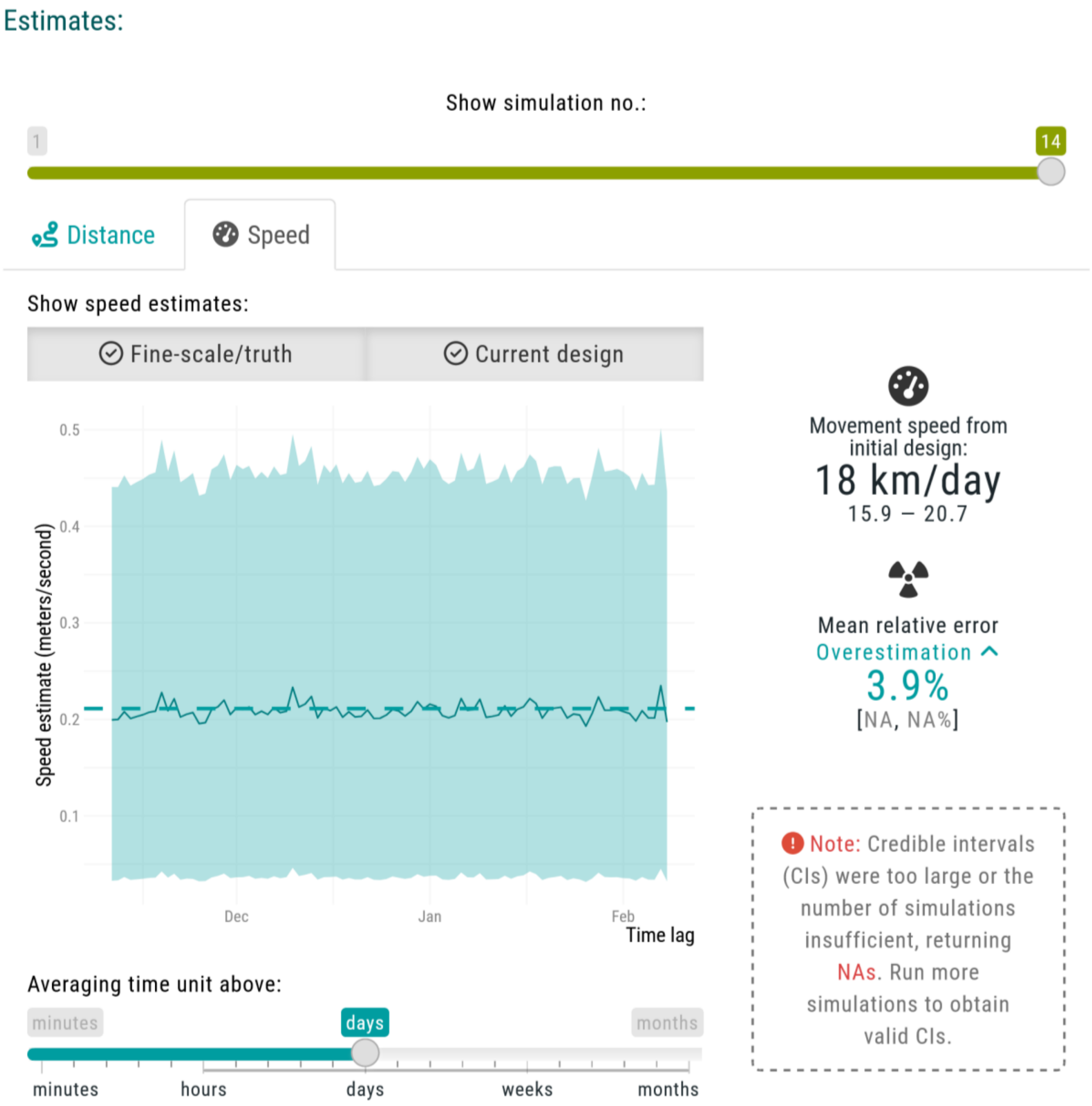


Next, in the ‘`Meta-analyses`’ tab, we click the ‘`Run meta-analyses`’ button to obtain group- and population-level inferences. Once completed, a new box on the left indicates that, on average, the **population-level home range area** is underestimated by `-22%` (`-35.2`, `-8.0%`), and that the **population-level movement speed** is overestimated by `6.2%` (`-1.3`, `14.1%`). In addition, two plots are generated: first with the **individual estimates of home range areas**, and then the **population-level estimates of home range areas**.

The **first plot** illustrates the relative error in **individual-level home range area estimates** by displaying the estimated home range area (x-axis, in km$^2$) for each group (y-axis), along with the associated 95% confidence intervals. The black square represents the population-level estimate (mean across all individuals) with its corresponding 95% confidence interval. The vertical solid lines indicate the expected true value for the inputted species parameters.

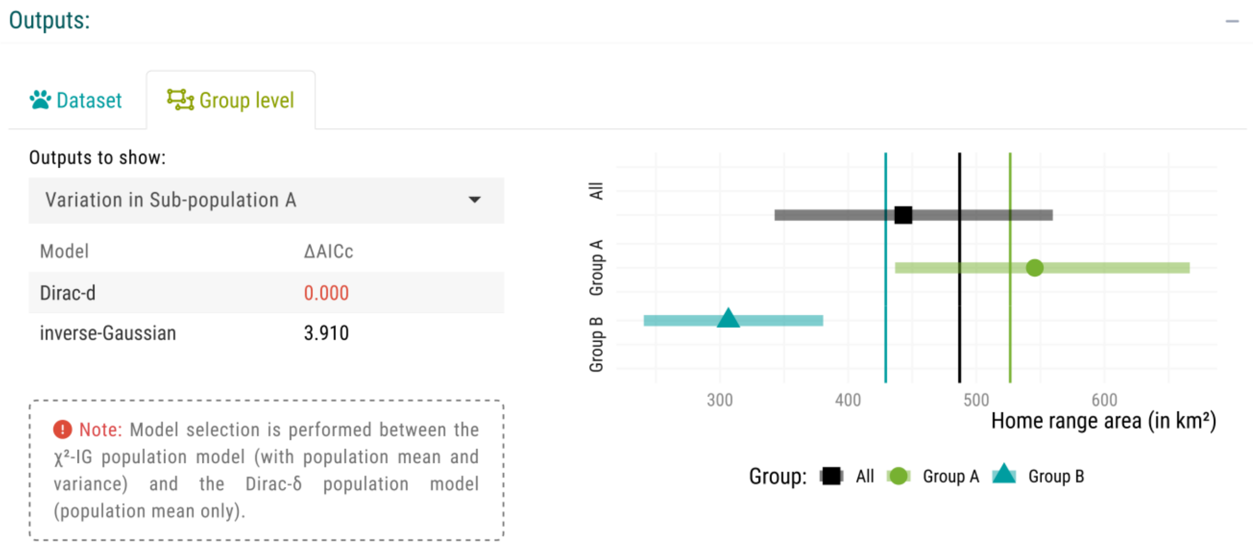


The **second plot** illustrates how an increasing number of individuals (from a *population* sample size of 3 to a maximum of 7, as the **total number of tags** was divided between the two groups) affects our **population-level mean estimates**. Each point represents the mean relative error (%) of this metric and its associated 95% confidence intervals, plotted against the number of tracked individuals. The dashed horizontal lines indicate a predefined error threshold of **±5%**. An accompanying table provides detailed numerical values, as well as whether a sub-population was detected at each *population* sample size.

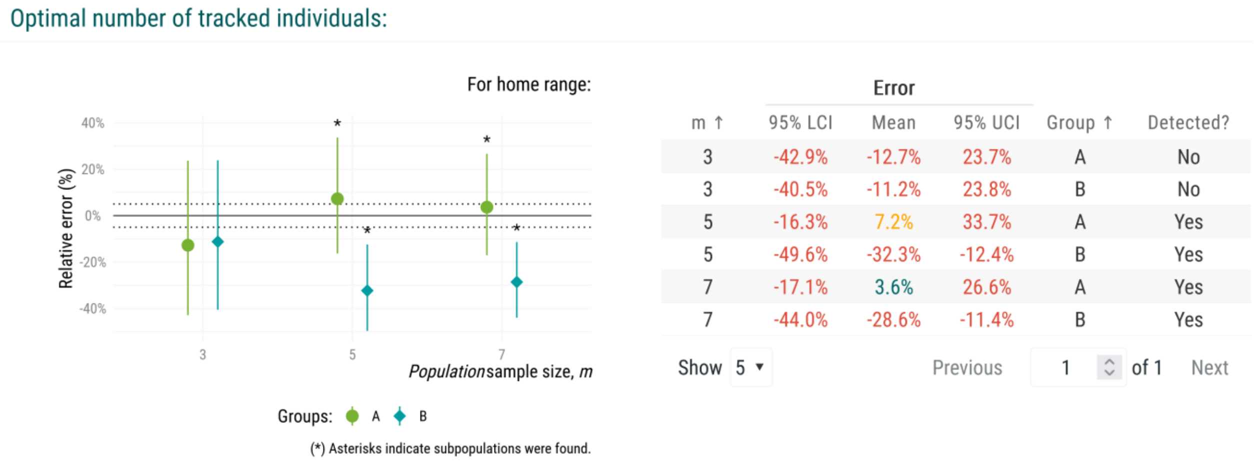


To verify these outputs, we can use a **resampling approach** to assess the spread of estimates, randomly reassigning individuals into new sets and rerunning the estimation of population-level mean estimates multiple times. This allows us to evaluate whether the observed mean estimates fall within the error threshold due to random chance, and assess the contribution of individual estimates. We set the resamples to 15 here, but higher values are recommended if simulating higher ***population* sample sizes**. Then, we click the **‘Resample’** button. The new plot illustrates how different sets of individuals shape the observed mean estimate across ***population* sample sizes**. The influence of individual variation remains noticeable, suggesting that increasing the number of tagged individuals could help stabilize the **mean home range area** estimate. We can also see that the mean relative error exceeds the acceptable threshold.

Optimal number of tracked individuals:

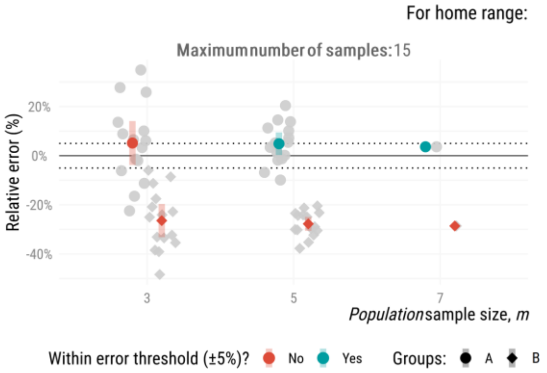


| m ↑ | Error 95% LCI | Error Mean | Error 95% UCI | Group ↑ | Detected? |
|---|---|---|---|---|---|
| 3 | -42.9% | -12.7% | 23.7% | A | No |
| 3 | -40.5% | -11.2% | 23.8% | B | No |
| 5 | -16.3% | 7.2% | 33.7% | A | Yes |
| 5 | -49.6% | -32.3% | -12.4% | B | Yes |
| 7 | -17.1% | 3.6% | 26.6% | A | Yes |
| 7 | -44.0% | -28.6% | -11.4% | B | Yes |

Show 5 ▾ Previous 1 of 1 Next

Finally, in the **‘Report’** tab, we can generate a final overview by clicking the **‘Build report’** button. This report highlights all the outputs from our current sampling schedule choice, informing us that 14 individuals is insufficient to obtain valid home range area or speed ratios, but it is sufficient to obtain a **population-level mean estimate** for **movement speed** within our error threshold of **±5%**.

*Population-level inferences report:*

Show outputs for:

Home range | Speed & distance

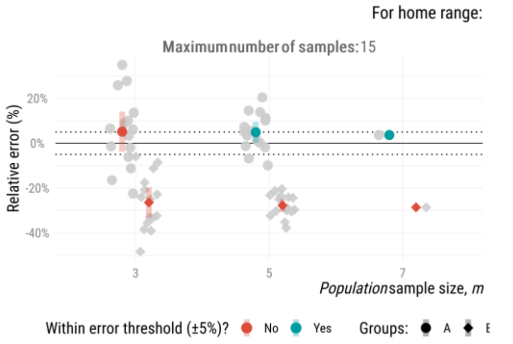


*Home range meta-analyses:*

You specified 14 tags, which is insufficient to produce a stable estimate of mean home range area within the error threshold of 5% The estimate is still shifting as resamples increase, meaning results cannot yet be considered reliable.

**The mean home range area based on 14 individuals (both groups) was overestimated by 0.6% [-19.5, 22.9]. Current study design is insufficient to accurately estimate home range area. Increase the number of tags to obtain a more reliable estimate of the population mean.**

There was insufficient evidence in the selected dataset to detect sub-populations. With the current parameters, sub-populations were detected (though with ΔAICc < 2). The home range area ratio (for group A/group B) did not overlap with one (ratio point estimate of 1.79:1). **The number of individuals appears insufficient to obtain valid home range area ratios.**

This workflow completed in 1 hour and 10 minutes on a laptop with an Intel Core Ultra 5 processor (1.6 GHz, 12 cores, 16 GB RAM) using parallel computation.

### **S1.2.** Determining the minimum number of units:

We can also evaluate study design iteratively by analyzing new subsets of individuals at a time, to determining the **minimum number of units** required to meet a predefined error threshold. Be sure to restart the application at this point to avoid any potential issues. For this workflow, we select a single **research target** (home range estimation), and mean estimate of sampled groups as the **analytical target**. Finally, for **deployment**, we choose "*I want to determine the minimum number of VHF/GPS tags*".

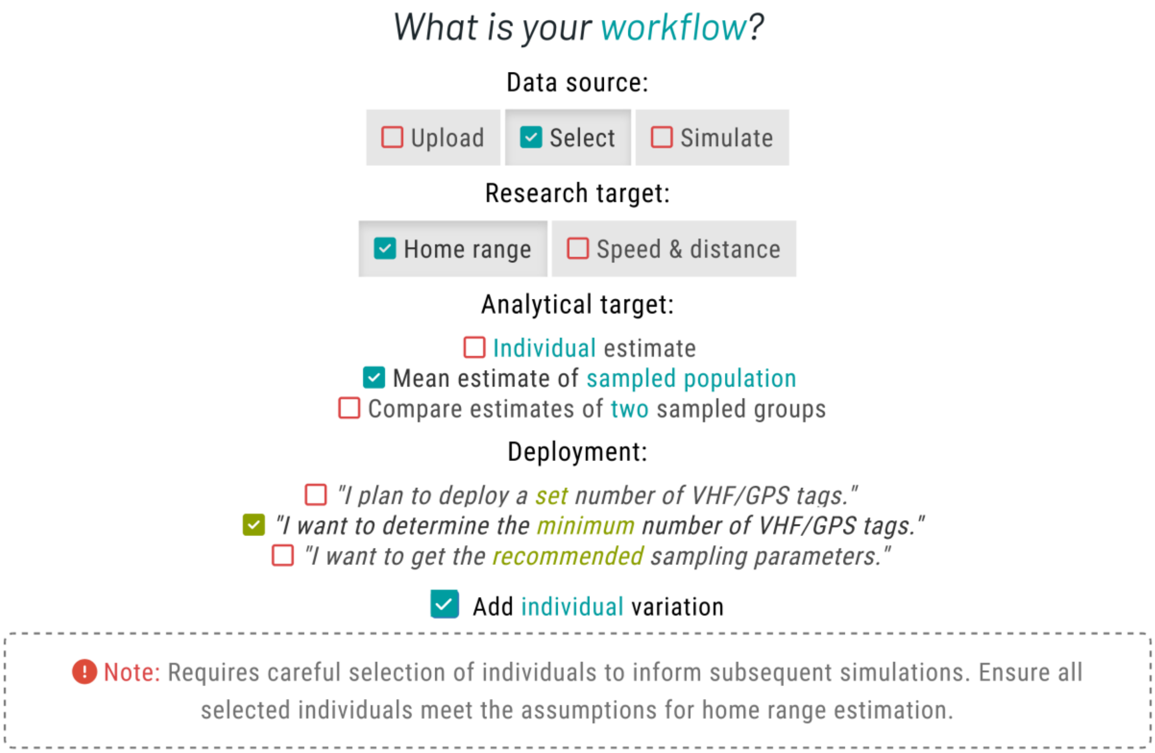


The only changes from the previous workflow are that we no longer need to set groups in the `Select data’` tab, and that, in the `‘Sampling design’` tab, we will evaluate a sampling schedule with a ***sampling duration*** of `2 years`, collecting one new location per day (***sampling interval*** of `1 day`). We will impose no further constraints.

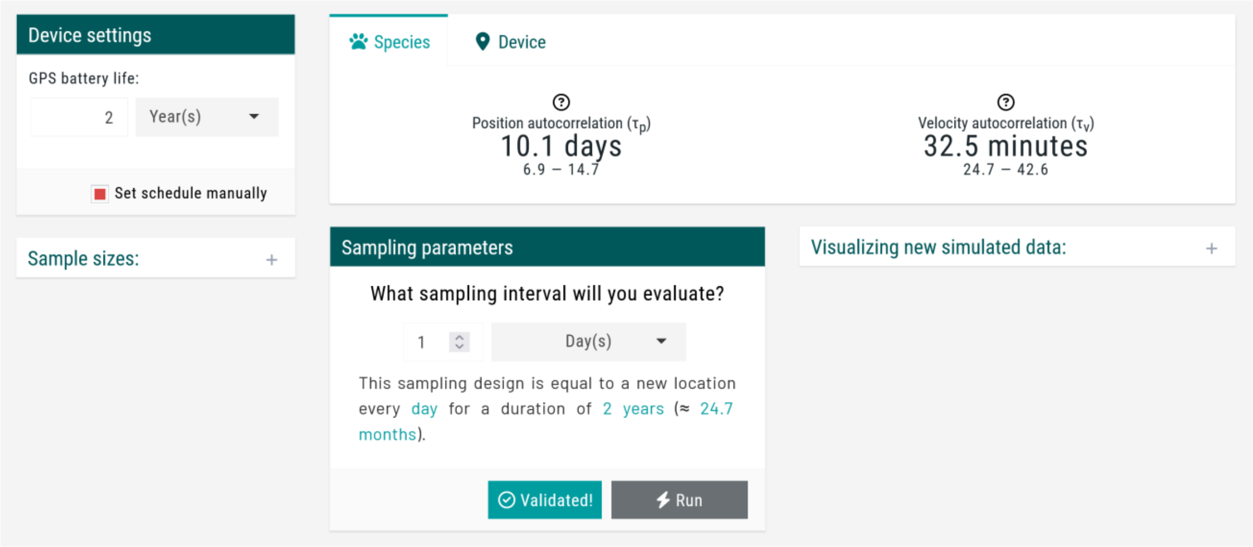


Once a confirmation message appears indicating a successful run up to this point, we proceed to the `'Home range'` tab to run an initial simulation. This step provides a preliminary estimate, allowing us to assess whether the setup and parameter choices align with our expectations. If necessary, we can refine the parameters before proceeding with further analyses.

Here, we will proceed with these sampling parameters and specify a maximum of `20` additional individuals (run in sets of `4`, each with `5` replicates), with an acceptable error threshold of `15%`. Simulations stop early when the mean estimate converges within the `±15%` threshold.

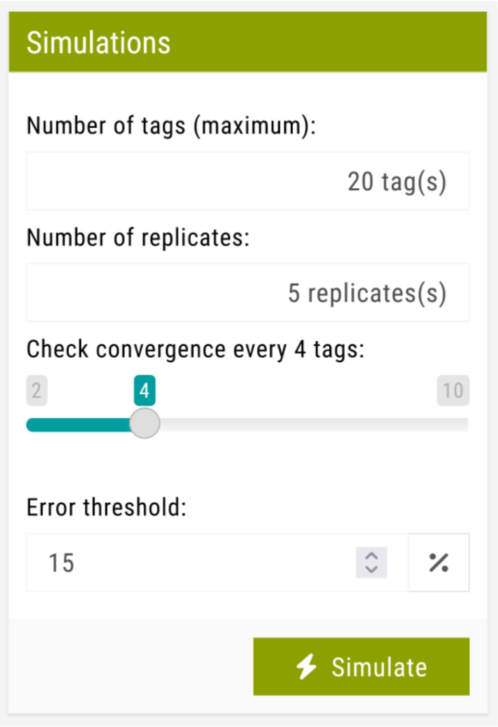

Once this is completed, a message will pop-up:

> **Minimum number of tags:**
>
> You specified a maximum of 20 tags. Under the current assumptions and an error threshold of 15%, a stable estimate of the population mean may be achieved by deploying 12 tags.
>
> If the **recommended number of tags** is close to (or equal to) the **maximum number of tags**, consider increasing the number of tags to reduce uncertainty. For a more detailed evaluation, explore the outputs in the Meta-analyses tab.

Then, we can evaluate these outputs in the `Meta-analyses` tab, where, results update automatically. Keep in mind that, due to the random nature of seed generation and the inclusion of individual variation, values may vary between runs. For our case, on average, the **population-level home range area** is underestimated by `-6.1%` (`-9.0`, `-3.3%`) at the ***population* sample size** of 12 individuals (when the error threshold was reached).

Optimal number of tracked individuals:

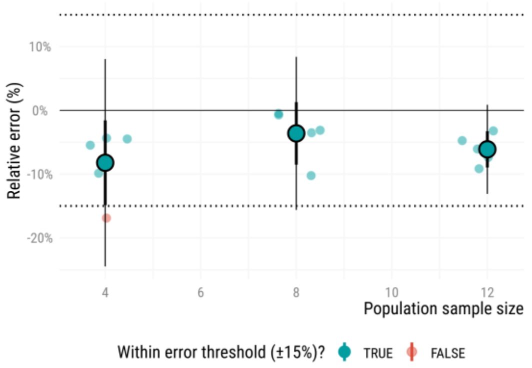


| | Error | | | | |
|---|---|---|---|---|---|
| m ↑ | 95% LCI | Mean | 95% UCI | Group ↑ | Detected? |
| 4 | -14.9% | -8.2% | -1.6% | All | 0.0% |
| 8 | -8.5% | -3.6% | 1.3% | All | 0.0% |
| 12 | -9.0% | -6.1% | -3.3% | All | 0.0% |

Replicate | 5 replicate(s)

Finally, in the `'Report'` tab, we can once again generate a final overview by clicking the `'Build report'` button:

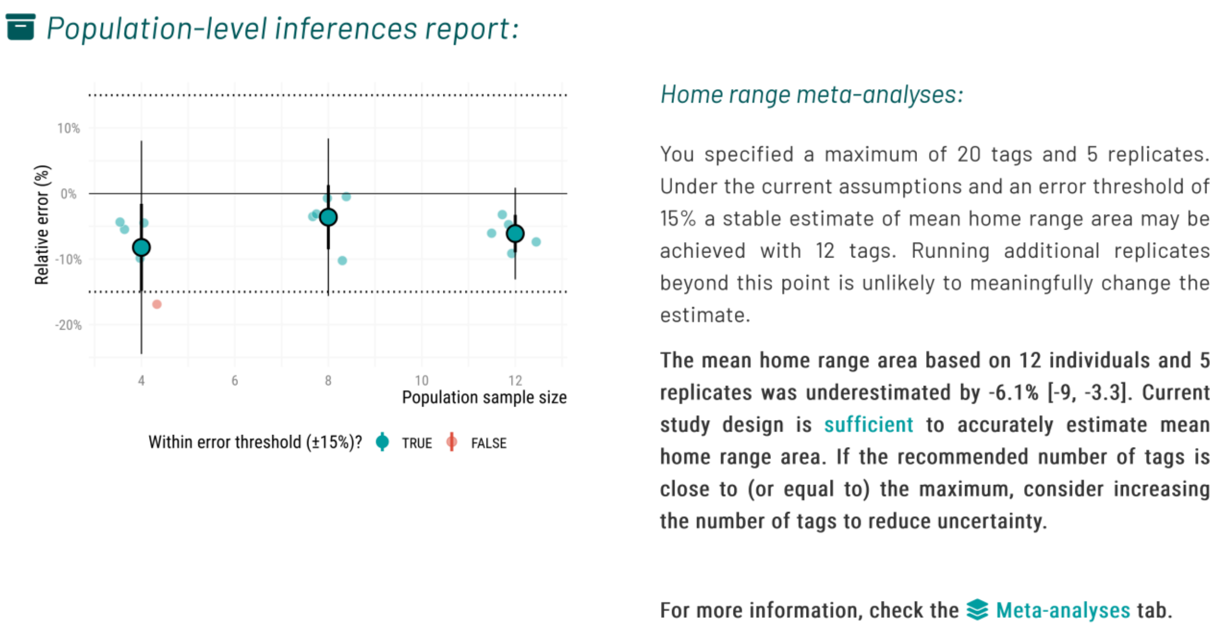


This workflow was completed in under 30 minutes on a laptop with an Intel Core Ultra 5 processor (1.6 GHz, 12 cores, 16 GB RAM) using parallel computation.
