## Supplementary material for "Too few, too many, or just right? Optimizing sample sizes for population-level inferences in animal tracking projects": Suplementary File S3

### **S3.** Single combination plots

#### Home range estimation

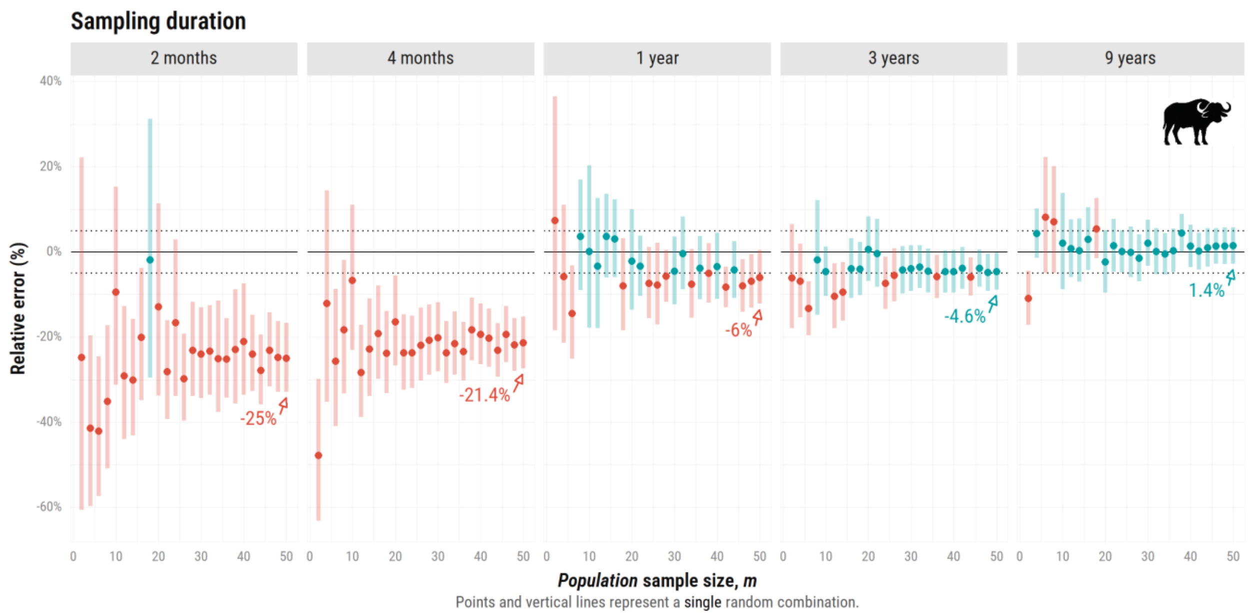


**Figure S3.1. Relative error (%) in home range estimates (AKDE) as a function of *population* sample size ($m$), ranging from 2 to 50 simulated African buffalos (*Syncerus caffer*).** Each vertical facet corresponds to one of five sampling durations: 2 months, 4 months, 1 year, 3 years, and 9 years. The plot shows mean estimates based on a **single** combination of individuals per $m$ (no resampling); estimates in blue are within the error threshold and in red if estimates fall outside of it. Horizontal dotted lines mark the ±5% error threshold.

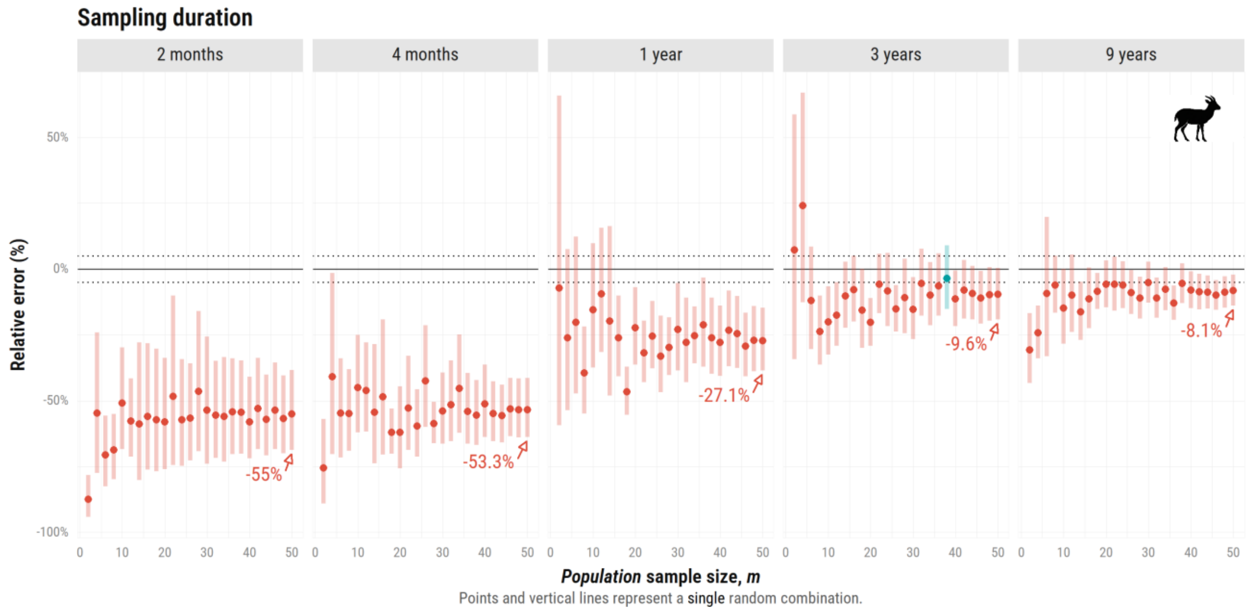


**Figure S3.2. Relative error (%) in home range estimates (AKDE) as a function of *population* sample size ($m$), ranging from 2 to 50 simulated Mongolian gazelles (*Procapra gutturosa*).** Each vertical facet corresponds to one of five sampling durations: 2 months, 4 months, 1 year, 3 years, and 9 years. The plot shows mean estimates based on a **single** combination of individuals per $m$ (no resampling); estimates in blue are within the error threshold and in red if estimates fall outside of it. Horizontal dotted lines mark the ±5% error threshold.

### Speed & distance estimation

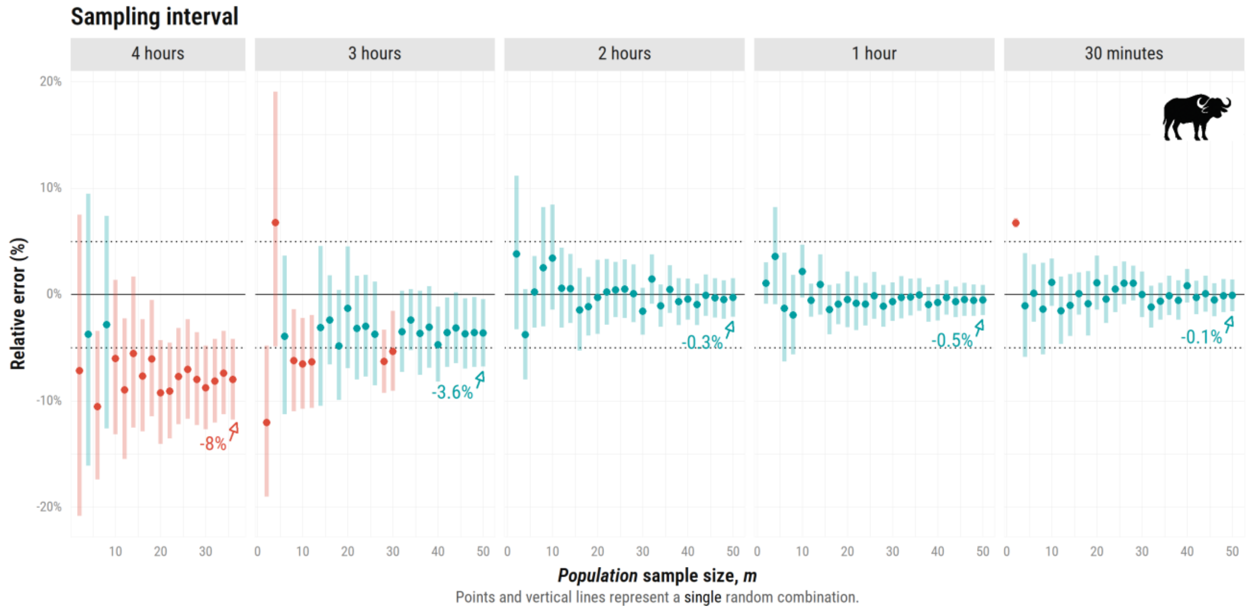


**Figure S3.3. Relative error (%) in speed estimates (CTSD) as a function of *population* sample size ($m$), ranging from 2 to 50 simulated African buffalos (*Syncerus caffer*).** Each vertical facet corresponds to one of five sampling intervals: 4 hours, 3 hours, 2 hours, 1 hour, and 30 minutes. The plot shows mean estimates based on a single combination of individuals per $m$ (no resampling); estimates in blue are within the error threshold and in red if estimates fall outside of it. Horizontal dotted lines mark the ±5% error threshold.

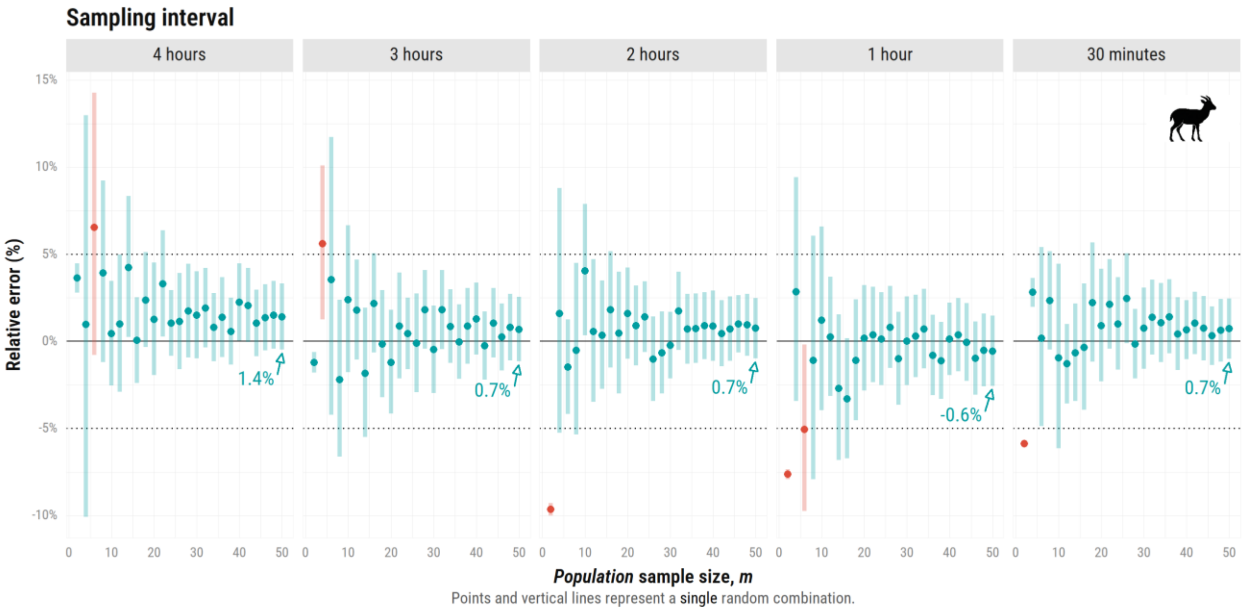


**Figure S3.4. Relative error (%) in speed estimates (CTSD) as a function of *population* sample size ($m$), ranging from 2 to 50 simulated Mongolian Gazelles (*Procapra gutturosa*).** Each vertical facet corresponds to one of five sampling intervals: 4 hours, 3 hours, 2 hours, 1 hour, and 30 minutes. The plot shows mean estimates based on a **single** combination of individuals per $m$ (no resampling); estimates in blue are within the error threshold and in red if estimates fall outside of it. Horizontal dotted lines mark the ±5% error threshold.
