## Supplementary material for "Too few, too many, or just right? Optimizing sample sizes for population-level inferences in animal tracking projects": Suplementary File S4

#### **S4.** Leave-one out approach

##### Home range estimation

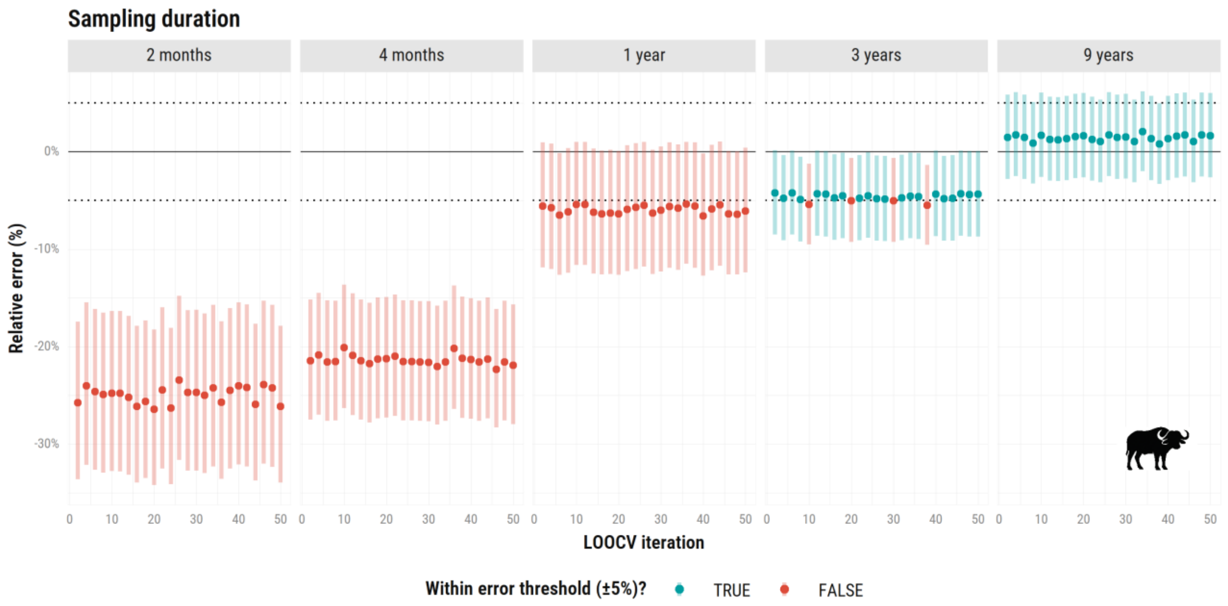


**Figure S4.1. Relative error (%) in home range area estimates across different sampling durations for simulated African buffalos (*Syncerus caffer*) for a leave-one-out approach.** Points represent the mean relative error (%) for one of 50 iterations (excluding a different individual each time), with its 95% confidence intervals. Horizontal dotted lines indicate the ±5% error threshold. Blue points fall within this threshold; red points fall outside of it.

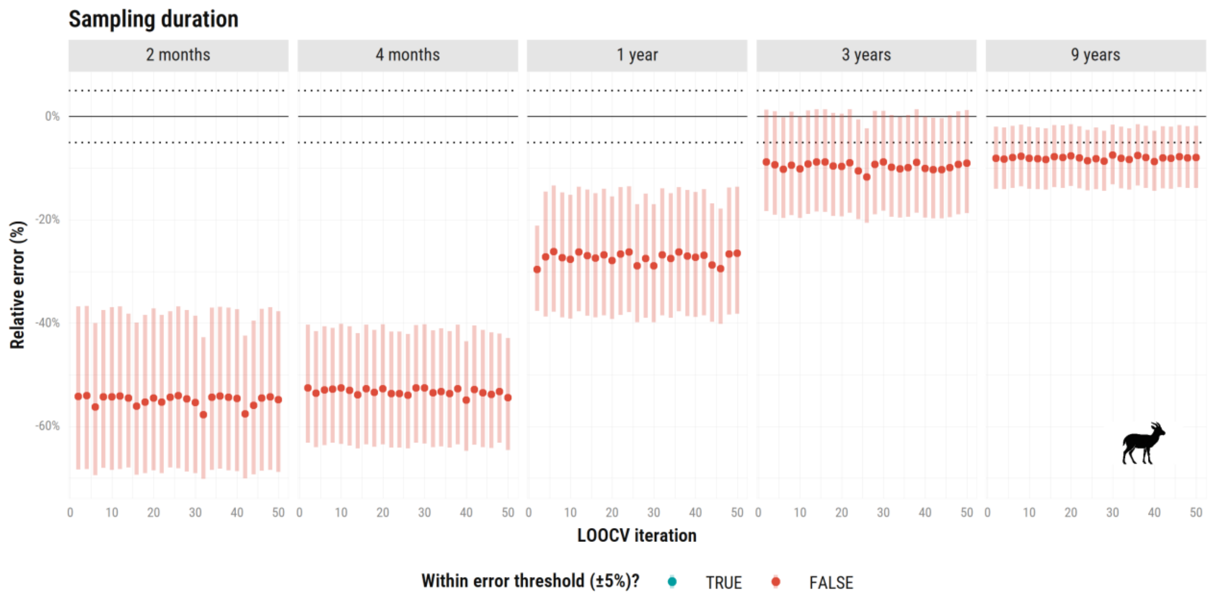


**Figure S4.2. Relative error (%) in home range area estimates across different sampling durations for simulated Mongolian gazelles (*Procapra gutturosa*) for a leave-one-out approach.** Points represent the mean relative error (%) for one of 50 iterations (excluding a different individual each time), with its 95% confidence intervals. Horizontal dotted lines indicate the ±5% error threshold. Blue points fall within this threshold; red points fall outside of it.

### Speed & distance estimation

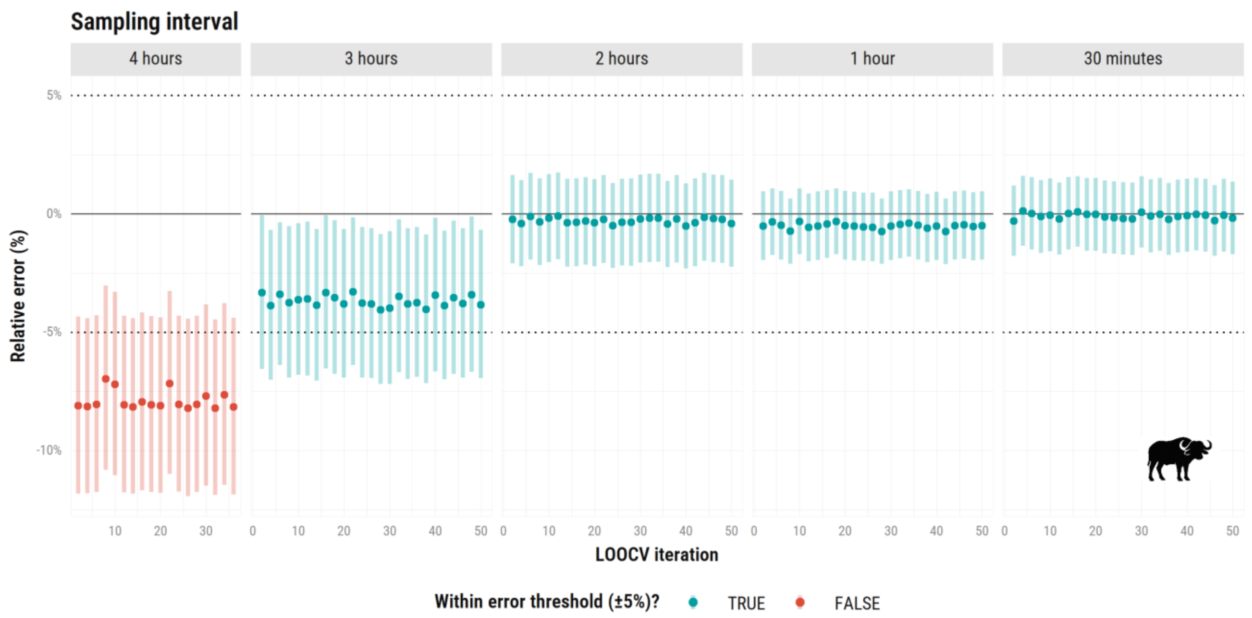


**Figure S4.3. Relative error (%) in speed & distance estimates across different sampling intervals for simulated African buffalos (*Syncerus caffer*) for a leave-one-out approach.** Points represent the mean relative error (%) for one of 50 iterations (excluding a different individual each time), with its 95% confidence intervals. Horizontal dotted lines indicate the ±5% error threshold. Blue points fall within this threshold; red points fall outside of it.

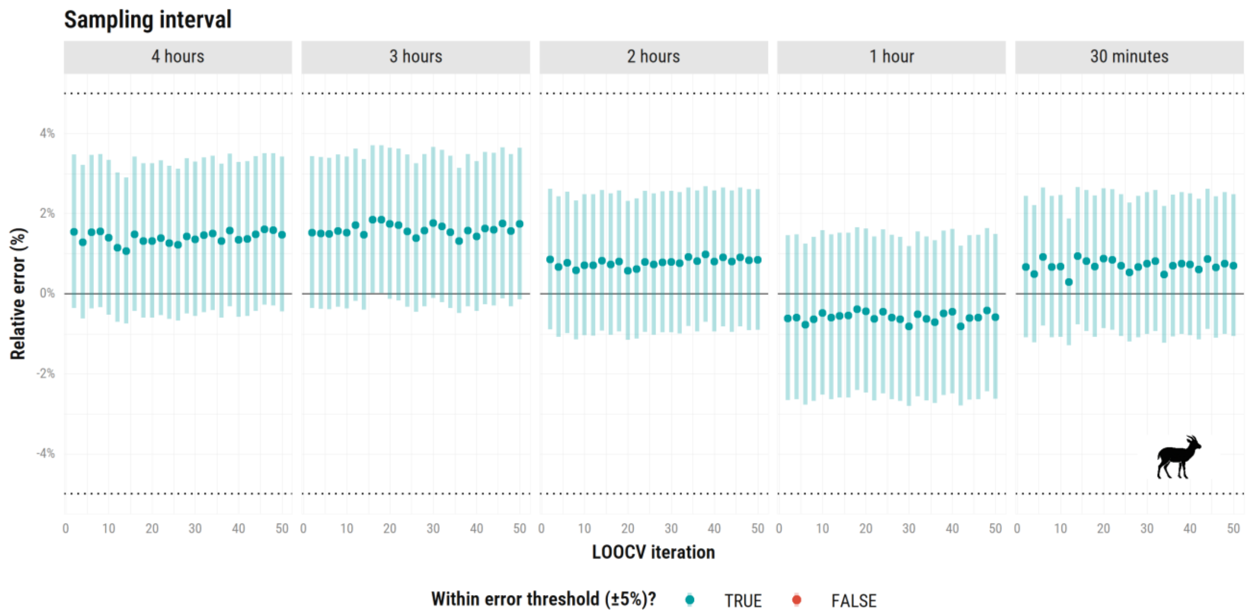


**Figure S4.4. Relative error (%) in speed & distance estimates across different sampling intervals for simulated Mongolian gazelles (*Procapra gutturosa*) for a leave-one-out approach.** Points represent the mean relative error (%) for one of 50 iterations (excluding a different individual each time), with its 95% confidence intervals. Horizontal dotted lines indicate the ±5% error threshold. Blue points fall within this threshold; red points fall outside of it.
